# Optimization of AsCas12a for combinatorial genetic screens in human cells

**DOI:** 10.1101/747170

**Authors:** Kendall R Sanson, Peter C DeWeirdt, Annabel K Sangree, Ruth E Hanna, Mudra Hegde, Teng Teng, Samantha M Borys, Christine Strand, J Keith Joung, Benjamin P Kleinstiver, Xuewen Pan, Alan Huang, John G Doench

**Affiliations:** Genetic Perturbation Platform, Broad Institute of MIT and Harvard, Cambridge, MA, USA; Tango Therapeutics, 100 Binney St., Cambridge, MA; Molecular Pathology Unit, Massachusetts General Hospital, Charlestown, MA, USA; Center for Cancer Research, Massachusetts General Hospital, Charlestown, MA, USA; Center for Computational and Integrative Biology, Massachusetts General Hospital, Charlestown, MA, USA; Department of Pathology, Harvard Medical School, Boston, MA, USA; Center for Genomic Medicine, Massachusetts General Hospital, Boston, MA 02114; Department of Pathology, Massachusetts General Hospital, Boston, MA 02114

## Abstract

Cas12a enzymes have attractive properties for scalable delivery of multiplexed perturbations, yet widespread usage has lagged behind Cas9-based strategies. Here we describe the optimization of Cas12a from *Acidaminococcus* (AsCas12a) for use in pooled genetic screens in human cells. By assaying the activity of thousands of guides, we confirm on-target design rules and extend them to an enhanced activity variant, enAsCas12a. We also develop the first comprehensive set of off-target rules for Cas12a, and demonstrate that we can predict and exclude promiscuous guides. Finally, to enable efficient higher-order multiplexing via lentiviral delivery, we screen thousands of direct repeat variants and identify 38 that outperform the wildtype sequence. We validate this optimized AsCas12a toolkit by targeting 12 synthetic lethal gene pairs with up to 400 guide pairs each, and demonstrate effective triple knockout via flow cytometry. These results establish AsCas12a as a robust system for combinatorial applications of CRISPR technology.

## INTRODUCTION

CRISPR technology has injected new life into the field of functional genomics, with its robust on-target activity, acceptable off-target profile, and a myriad of derivations that allow manipulation of the genome beyond genetic knockout^1^. Coupling CRISPR technology to highly parallel methods to write and read DNA, as well as viral technologies that enable delivery to nearly any cell type of interest, has enabled pooled genetic screens across diverse areas of biology^2–4^.

For individual perturbations, the Cas9 enzyme from *Streptococcus pyogenes* (SpCas9) was one of the first to be characterized^5^ and has seen the most subsequent development; it remains the tool of choice for most screening experiments. One use-case where SpCas9 is less optimal, however, is in the ability to multiplex, that is, the delivery of more than one guide RNA into cells using the same strategies compatible with pooled genetic screens. The need to have individual expression cassettes for each additional guide RNA requires multi-step cloning to generate libraries and customized sequencing readouts^6^; further, such arrangements are prone to recombination^7^ and uncoupling^8,9^ when made into lentivirus and retrieved by PCR. Thus, there is a need for a more robust system for perturbing multiple genes at the same time.

The Cas12a family of CRISPR enzymes (previously known as Cpf1) potentially offers a better approach for multiplexing, because an array of guide RNAs can be expressed from a single transcript, separated only by a 20 nucleotide direct repeat (DR) sequence, and Cas12a is able to both process individual guides out of the array and then execute target DNA cleavage^10^. Several Cas12a orthologs have shown activity in human cells, including variants derived from *Acidaminococcus* (AsCas12a) and *Lachnospiraceae* (LbCas12a)^10^, and the nucleotide preferences for both of these orthologs have been broadly assessed^11,12^. Further, LbCas12a has been developed for CRISPRa approaches^13^ and base editing^14^, whereas AsCas12a has undergone protein engineering to increase efficacy and is thus another option for CRISPRa and base editing approaches^15^. Other Cas12a orthologs have also seen continued development^16,17^. To-date, however, only one small scale genetic screen has taken advantage of the multiplexing ability of Cas12a, but it relied on stringent positive selection^18^, which makes assessment of efficiency difficult. Further, even when delivered by transient transfection, which is generally a poor surrogate for the use of single copy lentivirus needed for pooled screens^19^, proof-of-concept multiplexing assays have demonstrated poor efficiency, with only a small fraction of cells showing all intended edits^20^.

Here we optimize the AsCas12a system for pooled genetic screens in human cells. We test several variants of AsCas12a to select a construct with high efficiency, and we validate on-target sequence rules for selecting guides. Because avoidance of promiscuous guides is critical for library design, we screened mismatched guides to derive rules for avoiding off-target cutting. Additionally, we develop alternative direct repeat sequences to avoid recombination in the multiplexed array. Finally, we validate these optimized components by conducting a dual knockout screen for synthetic lethality as well as a triple knockout flow cytometry assay. In sum, we present a complete set of experimental and computational tools to enable the effective use of AsCas12a in combinatorial screening applications.

## RESULTS

### Optimization of the AsCas12a protein

To assess the potential of AsCas12a for screening purposes, we first acquired a lentiviral vector described in the literature^11^, which has a single nuclear localization sequence (**Fig. 1a**), referred to here as 1xNLS-Cas12a. We also constructed a second lentiviral vector that includes two NLSs (2xNLS-Cas12a). This served as a template for a third vector, into which we introduced the amino acid point mutations to generate enCas12a, a recently-described variant of AsCas12a with increased activity and an expanded range of PAM sequences^15^.

**Figure 1.**
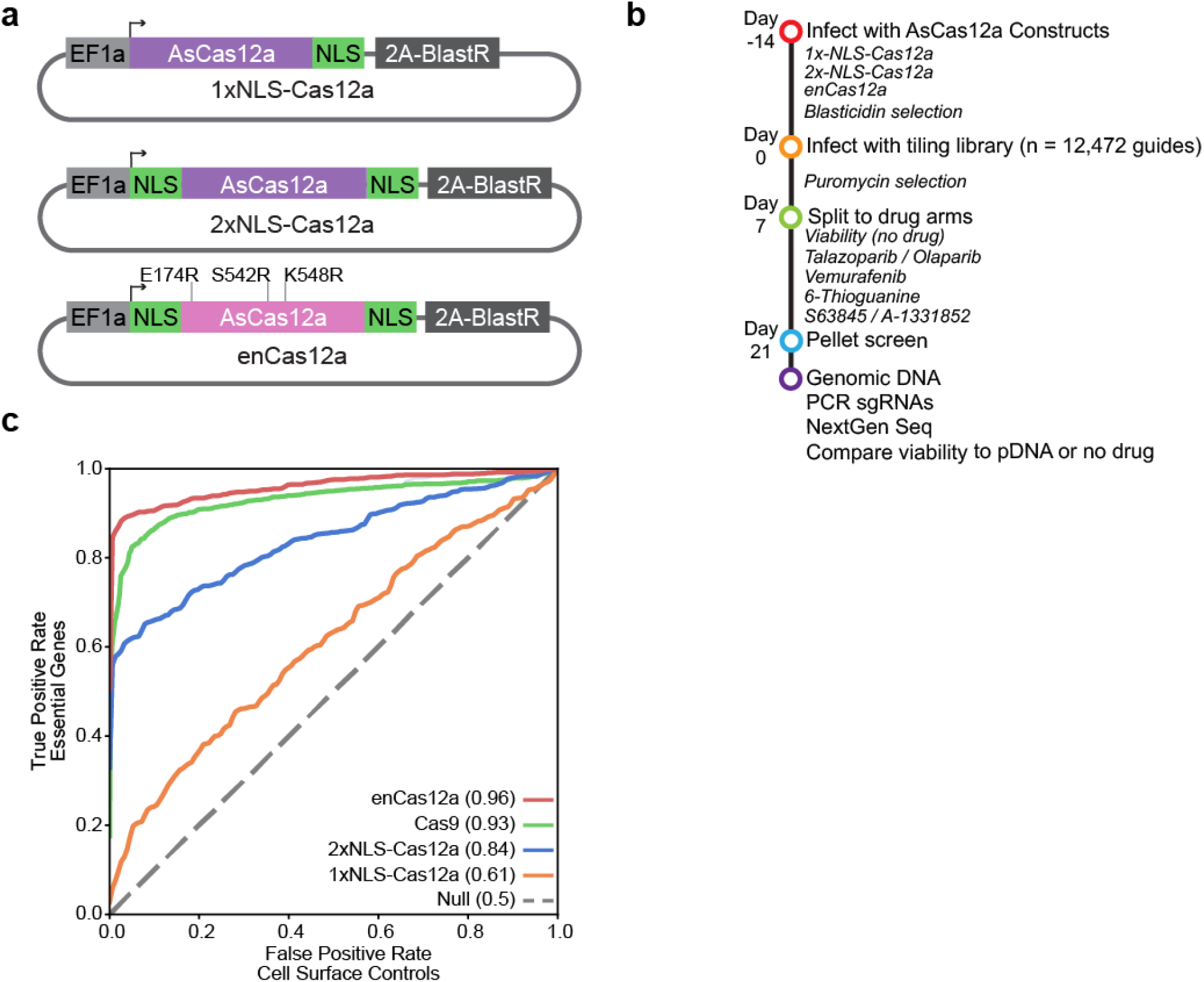
Optimization of AsCas12a for use in pooled screening. (**a**) Vector maps for Cas12a constructs. Point mutations for enCas12a are indicated. (**b**) Timeline for executing tiling screens. (**c**) Precision-recall curves for guides targeting essential and control genes for Cas12a and SpCas9, using viability data in A375 cells (n = 1146 and 2468 for essential guides for Cas12a and Cas9, respectively, and 153 and 673 for control guides). The area under the curve for each enzyme is noted in parentheses.

We synthesized a tiling library of all possible guides ranging between 20 and 23 nucleotides in length targeting TTTN PAMs across a set of 43 genes. This library included pan essential genes^21^, cell-specific lethal genes, genes with well-characterized interactions with specific small-molecules, and cell surface genes, as well as all possible guides for the essential gene EEF2 regardless of PAM, for a total of 12,472 unique guides (**Supplementary Table 1**). We also synthesized an analogous version of this library for use with SpCas9 (NGG PAM), to enable direct comparison of activity. The AsCas12a library was cloned into a modified version of lentiGuide, pRDA_052, which expresses the guide from a U6 promoter and confers puromycin resistance. Positive and negative selection screens were conducted over a three week time course in duplicate (**Fig. 1b**), as we have done previously to assess activity of SaCas9^22^ and SpCas9^23^ via growth assays. During the course of this study, the AsCas12a tiling library was screened across five different cell lines and six small molecule treatments (**Supplementary Table 2**, **Supplementary Data 1**).

We first examined the viability data from A375 cells, as this assay was conducted with all three AsCas12a constructs as well as SpCas9. For each of the AsCas12a variants, we observed similar performance across all lengths of guide RNAs (**Supplementary Fig. 1a**). Further, we saw the highest activity at TTTA PAM sites followed closely by TTTC and TTTG, and the lowest activity levels at TTTT sites for all variants (**Supplementary Fig. 1b**), which has been observed previously^11^. Defining the 5^th^ percentile of guides targeting control genes as a cutoff for activity, we saw 57% of enCas12a guides targeting a TTTT site were active compared with 7% of 2xNLS-Cas12a and 7% of 1xNLS-Cas12a, demonstrating the ability of enCas12a to target previously inaccessible sites^11,15^. Comparing across cell lines (**Supplementary Fig. 1c**) we saw that guides that were effective in one tended to be effective in another.

To compare the efficacy of AsCas12a to SpCas9, we performed a precision recall analysis with the A375 viability data (excluding guides with TTTT PAMs for AsCas12a), defining guides targeting essential genes as true positives and guides targeting control genes as false positives, and calculating an area under curve (AUC) (**Fig. 1c**). SpCas9 performed well, with an AUC of 0.93, whereas the 1xNLS-Cas12a construct performed the worst with an AUC of 0.61. However, the additional NLS site improved performance, as the AUC with 2xNLS-Cas12a increased to 0.84, consistent with recent observations^24^. Finally, enCas12 showed the most robust performance, comparable to SpCas9, with an AUC 0.96. Thus, we chose to move forward with 2x-NLS-Cas12a and the enCas12a variant thereof for further experiments.

### Evaluation of on-target scoring criteria

We used data from the tiling screens to evaluate a published deep learning model, referred to as Seq-DeepCpf1^12^. This model was trained using 15,000 insertion and deletion (indel) frequencies, generated by integrating guides and target sequences into the same cassette and read out with deep sequencing^11^. We compared this model to a gradient boosted (GB) model trained on the same set of indel frequencies, as well as two GB models trained on 2xNLS-Cas12a and enCas12a tiling data (**Fig. 2a**). We fit the GB models using the same framework that we used for Cas9 ruleset development^22,23^, and held out a test set for all three sources of data (**Supplementary Fig. 2a**). For each test set we evaluated guides targeting TTTN and TTTV PAMs separately, due to the relatively low activity of TTTT PAM targets. We observed that Seq-DeepCpf1 performed well across test sets (**Fig. 2b**). Further, the GB model trained on the indel frequencies performed similarly to Seq-DeepCpf1, illustrating the high quality of this training dataset. We saw that Seq-DeepCpf1 and the 2xNLS-Cas12a GB model made similar predictions on the 2xNLS-Cas12a test set (**Fig. 2c**), despite using orthogonal training data, suggesting that these models learned similar guide features. These results confirm the performance and generalizability of Seq-DeepCpf1.

**Figure 2.**
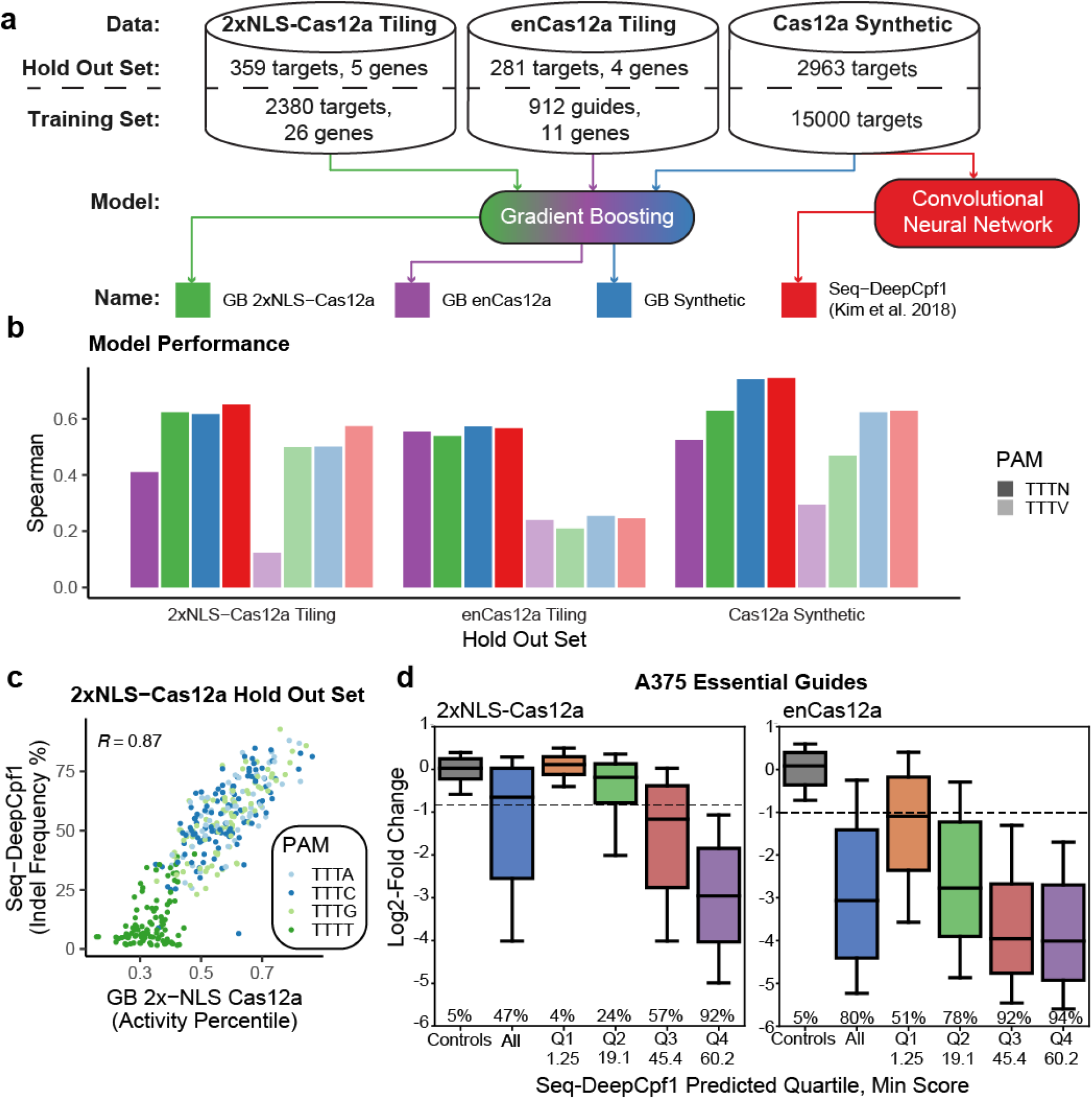
Evaluation of AsCas12a on-target predictions. (**a**) Summary of models under consideration. (**b**) Model performance on held out test sets. Spearman correlation shown for TTTN (dark) and TTTV only (light) target sites. Hold-out set sizes are noted in (a). (**c**) Predictions of GB 2xNLS-Cas12a and Seq-DeepCpf1 on the 2xNLS-Cas12a hold out set. Spearman correlation is included. (**d**) Activity of essential guides in A375 cells binned by predicted quartile for 2xNLS-Cas12a and enCas12a (n = 1589 essential guides, 231 control guides) Boxes represent the 25th, 50th and 75th percentiles, whiskers show 10th and 90th percentiles. Minimum score for each quartile is indicated.

To determine if the underlying nucleotide preferences for 2xNLS-Cas12a and enCas12a were similar, we compared the learned nucleotide importances of the two GB models using *in-silico* saliency analysis^25^ (see methods). The models agreed on the most extreme features (**Supplementary Fig. 2b**), but also showed some variation for smaller features (**Supplementary Fig. 2c, d**), suggesting that the underlying affinities for the two AsCas12a variants are similar but not identical. This means that a model trained with wildtype AsCas12a data can reliably identify the most important nucleotide features for enCas12a as well. Thus, we opted to use Seq-DeepCpf1 to generate on-target scores for 2xNLS-Cas12a and enCas12a moving forward.

To understand how to best use Seq-DeepCpf1 scores to select active guides, we focused on guides targeting essential genes in the tiling library. As expected, when we binned guides by Seq-DeepCpf1 score we saw agreement between predicted quartiles and observed activity for both 2xNLS-Cas12a and enCas12a (**Fig. 2d**). Using the 5^th^ percentile of cell surface control guides as an activity cutoff, we saw 92% and 94% of guides were active for 2xNLS-Cas12a and enCas12a in the quartile with the highest prediction scores, compared with 47% and 80% for all guides, respectively. Furthermore, when we selected the top half of essential and non-essential guides by Seq-DeepCpf1 score from the 2x-NLS and enCas12a tiling screens we saw the AUC increase to 0.94 and 0.97, respectively (**Supplementary Fig. 3**). Thus, applying Seq-DeepCpf1 scores increased the efficacy of guides for both constructs.

One major advantage of enCas12a is its ability to target a broader range of PAM sites, as the PAM for wildtype AsCas12a, TTTV, will occur only once every 96 nts on average, compared to NGG for SpCas9, which will occur once every 8 nts. The expanded PAMs of enCas12a were originally classified into three tiers^15^, with the first being the most active. To evaluate these alternative PAMs we focused on the set of EEF2 guides in the tiling library that targeted non-canonical PAMs (non-TTTN). These controls covered 249 out of 264 possible PAMs with at least one guide. In agreement with previous results, we saw a relationship between the assigned tier and the measured activity (**Supplementary Fig. 4a**). Notably, the guides targeting TTTN (62% active) showed similar efficacy to the other tier 1 PAMs (68% active).

Because none of these alternative PAMs were included in the training set for Seq-DeepCpf1, we reasoned that on-target activity predictions would be hindered if the model were provided with a non-TTTN PAM. We hypothesized that on-target predictions may improve by modifying the target sequences *in silico* to include a canonical PAM (**Supplementary Fig. 4b**). We saw that substituting TTTC for the actual, non-canonical PAM improved on-target predictions when compared to unaltered sequences for all three PAM tiers (**Supplementary Fig. 4c**), identifying highly active guides targeting non-canonical PAMs (**Supplementary Fig. 4d**). For the quartile of guides that had the highest Seq-DeepCpf1 scores, we saw that 93% of non-canonical tier 1 guides were active, which is comparable to the activity of guides targeting canonical PAMs. Since enCas12a also had increased activity at TTTT PAM sites, we performed a similar analysis focusing on essential guides targeting this PAM. We found that predictions also improved by *in silico* modification of the PAM (**Supplementary Fig. 4e**). Thus, at least until more data can be acquired to properly train an activity model for enCas12a at non-canonical PAMs, modification of the input sequence to Seq-DeepCpf1 can be useful for selecting active guides. The ability to select active guides from the expanded set of Tier 1 PAMs increases the frequency of targetable sites to approximately once every 7 nts, an important criterion for broad utility of AsCas12a.

### Off-target predictions for Cas12a

In order to assess the off-target tolerance of AsCas12a, we constructed a library of guides intentionally mismatched to their target. We selected 300 of the most active guides from the viability, vemurafenib, and 6-thioguanine screens with the tiling library (**Table 1**) in both A375 and Meljuso. We then designed every possible single and double nucleotide mismatch for these guides, and selected a random subset of the latter, resulting in a library with 300 perfect match guides, 19,512 single mismatch guides, 20,000 double mismatch guides, and finally 217 guides targeting cell surface genes as a set of negative controls (**Fig 3a**). We performed screens in duplicate in A375 cells with dropout, vemurafenib, and 6-thioguanine arms in cells stably expressing 2xNLS-Cas12a and enCas12a. In order to determine the fraction of guides that were active in this library we mapped the distribution of guides targeting essential genes in dropout assays and set an activity cutoff at the 5^th^ percentile of controls (**Fig 3b**). As expected, we found the perfect match guides to be highly active for both constructs, with 94% and 91% of guides scoring as active with 2xNLS-Cas12a and enCas12a, respectively. We observed that enCas12a showed a higher tolerance for mismatches, with 54% and 16% of guides active for single and double mismatch guides respectively, compared with 28% and 9% for 2xNLS-Cas12a. Thus, enCas12a shows more propensity for off-target activity, as initially described^15^.

**Figure 3.**
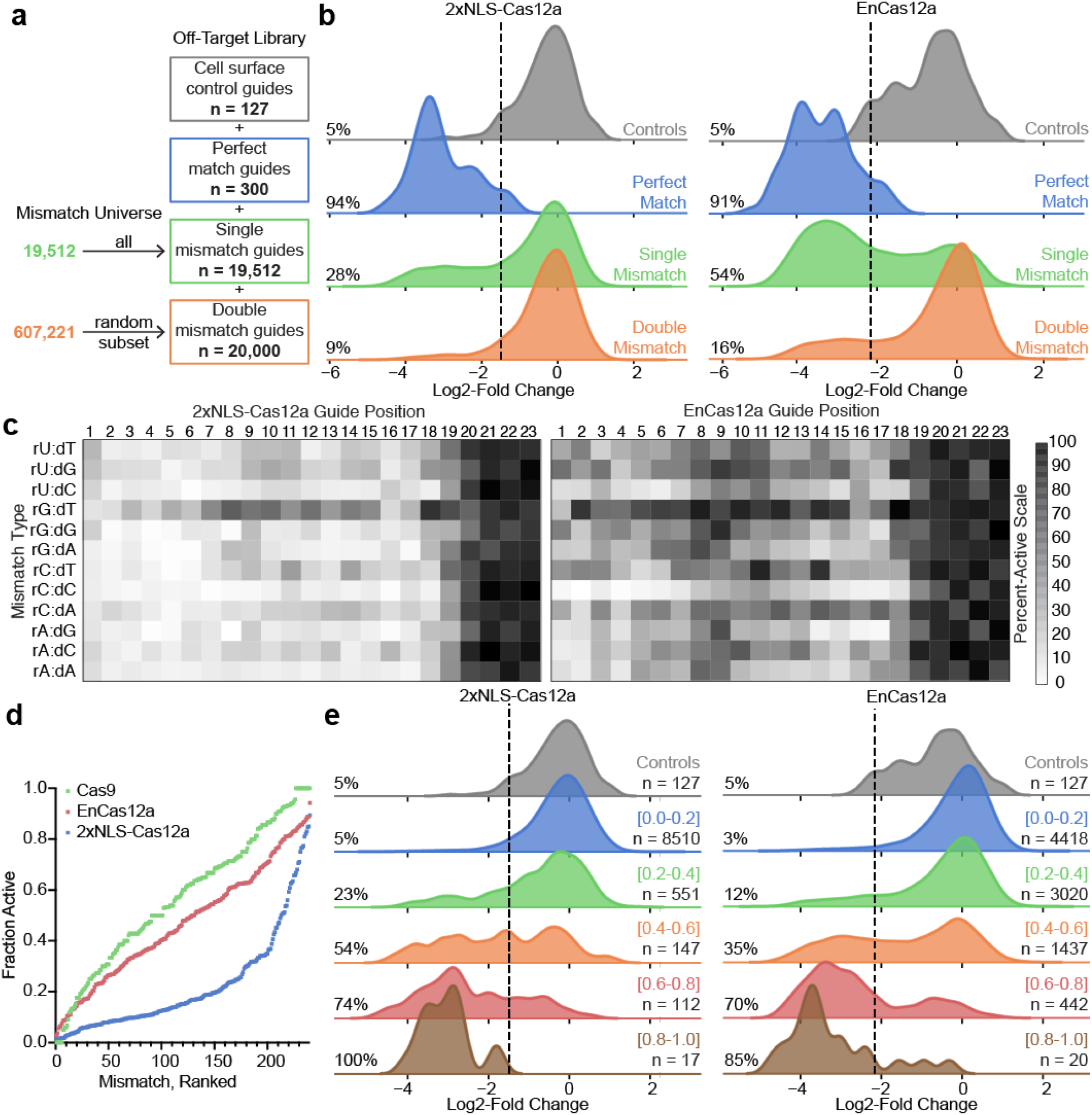
Prediction of off-target activity for AsCas12a (**a**) Schematic depicting off-target library construction and guide selection. (**b**) Density plot showing activity of guides in dropout screens targeting essential genes with zero, one and two mismatches. Line is displayed at the 5th percentile of guides targeting control genes, and used as a cutoff to determine the percent of guides active in each tier as reported. (**c**) Heat map displaying the fraction of guides active for each mismatch type at a given position in the guide. Guide position is numbered such that position 1 is PAM proximal. The fraction of active guides is reported from essential guides in dropout assays, vemurafenib resistance genes in vemurafenib assays, and HPRT1 guides in 6-thioguanine assays. Cutoff to determine active guides was determined by the 5th percentile of control genes in negative selection assays and the 95th percentile of control genes in positive selection assays. (**d**) Comparison of off-target activity of Cas9, 2xNLS-Cas12a, and enCas12a constructs. For each construct, the CFD scores of each position and nucleotide type were ranked, and the values plotted in ascending order. For Cas12a, only the first 20 nucleotides of the guide are used. Cas9 data are from a previous publication^23^. (**e**) Histograms displaying measured activity of double mismatch guides targeting essential genes in dropout scores binned by a prediction of activity, the Cutting Frequency Determination (CFD) score. The fraction activity determined by the fraction of guides that fell below the 5% flow control genes cutoff is reported for each CFD bin as well as the number of guides that fall into each bin.

We then calculated the fraction of active guides for each mismatch type and position to generate a cutting frequency determination (CFD) matrix, as we had done previously with SpCas9^23^. These matrices were similar across experimental conditions (**Supplementary Fig 5a**), so we merged the data to create a single CFD matrix for each Cas12a (**Fig. 3c**). When we compared the CFD values for 2x-NLS-Cas12a and enCas12a, we saw a monotonic relationship (**Supplementary Fig 5b**), indicating similar specificity preferences. For both constructs we saw a higher tolerance for mismatches at the PAM distal end of the guide, as well as for rG:dT mismatches, which are two trends observed previously with other techniques to examine off-target activity of Cas12a enzymes^26,27^, and that have also been seen with SpCas9^23,28^.

To compare CFD matrices between the two AsCas12a constructs and SpCas9, we used data from our previously published SpCas9 CFD matrix^23^. Since SpCas9 guides are designed as 20mers, we focused on the 20 most PAM-proximal nucleotides of AsCas12a guides. We then ranked each mismatch and position by activity and saw that SpCas9 and enCas12a had similar activity levels across their profiles, whereas 2xNLS-Cas12a was the least promiscuous (**Fig 3d**), consistent with previous examinations of the specificity of Cas12a enzymes by orthogonal techniques^26^. Thus, although enCas12a is more promiscuous than 2xNLS-Cas12a, its specificity is comparable to that of SpCas9, suggesting it is suitable for genetic screens.

Finally, we used the CFD matrix to predict the activity of double mismatch guides. To do this, we calculated a CFD score as the product of the activities of each individual mismatch, an approach that has also been used by others to identify problematic off-target sites with more than one mismatch for SpCas9^29,30^. To evaluate this model, we binned double mismatch guides targeting essential genes into predicted quartiles and plotted the distribution of measured log2-fold-change values in the dropout screens (**Fig 3e**). We saw more activity from guides with a larger CFD score, indicating that this model can help identify problematic multi-mismatch off-target sites when designing guides. Notably, the largest portion of double mismatch guides fell into the lowest quartile of CFD scores (**Fig. 3e**), which indicates that consideration of off-target activities does not overly restrict the set of guide RNAs available to target a gene of interest.

### Development of variant direct repeat sequences

When working with lentivirus, recombination of repetitive elements and shuffling of vectors at sites of homology can be a technical hurdle^7,9,22,31^. Effective multiplexing with Cas12a requires delivery of multiple direct repeats (DRs) within a single array, separated by only the 20 - 23 nt guide sequence. Thus, we sought to find variants of the DR sequence that would enable effective multiplexing, but would be less prone to potential recombination. We designed a library of 35,682 alternative DRs with up to 3 variable basepairs in the stem and 3 variable nucleotides in the single-stranded or loop regions. We also included two variations of control sequences: random sequences of 20 nucleotides; and transcription termination sequences, with a run of 6 thymidines.

To assay the efficacy of these DRs in a negative selection screen, we targeted BCL2L1 and MCL1, a known synthetic lethal pair^6,22,32^, such that only effective use of both guide RNAs in the same cell should lead to cell death. We cloned the library of DR sequences into two vectors, pRDA_127 which has a BCL2L1 guide in the first position and an MCL1 guide in the second position, and pRDA_128, which has the order of guides reversed (**Fig 4a**). We introduced the DR library into 2xNLS-Cas12a expressing Meljuso cells and cultured the population for 19 days before harvesting genomic DNA, PCR-amplifying the inserts, and quantitating the DR sequences (**Fig 4b**). Both pRDA_127 and pRDA_128 showed good reproducibility across replicates (**Supplementary Fig. 6a, b**). Because loss of BCL2L1 has a slight growth effect on its own^22^, we saw a shift in the control DR sequences that had a BCL2L1 guide in the first position followed by a truncated sequence, compared with an MCL1 guide in the first position (**Supplementary Fig. 6c, d**). From the 35,682 DR variants we tested, we identified 38 sequences which had a greater log2-fold change than the wildtype sequence across the two constructs (**Fig 4c**).

**Figure 4.**
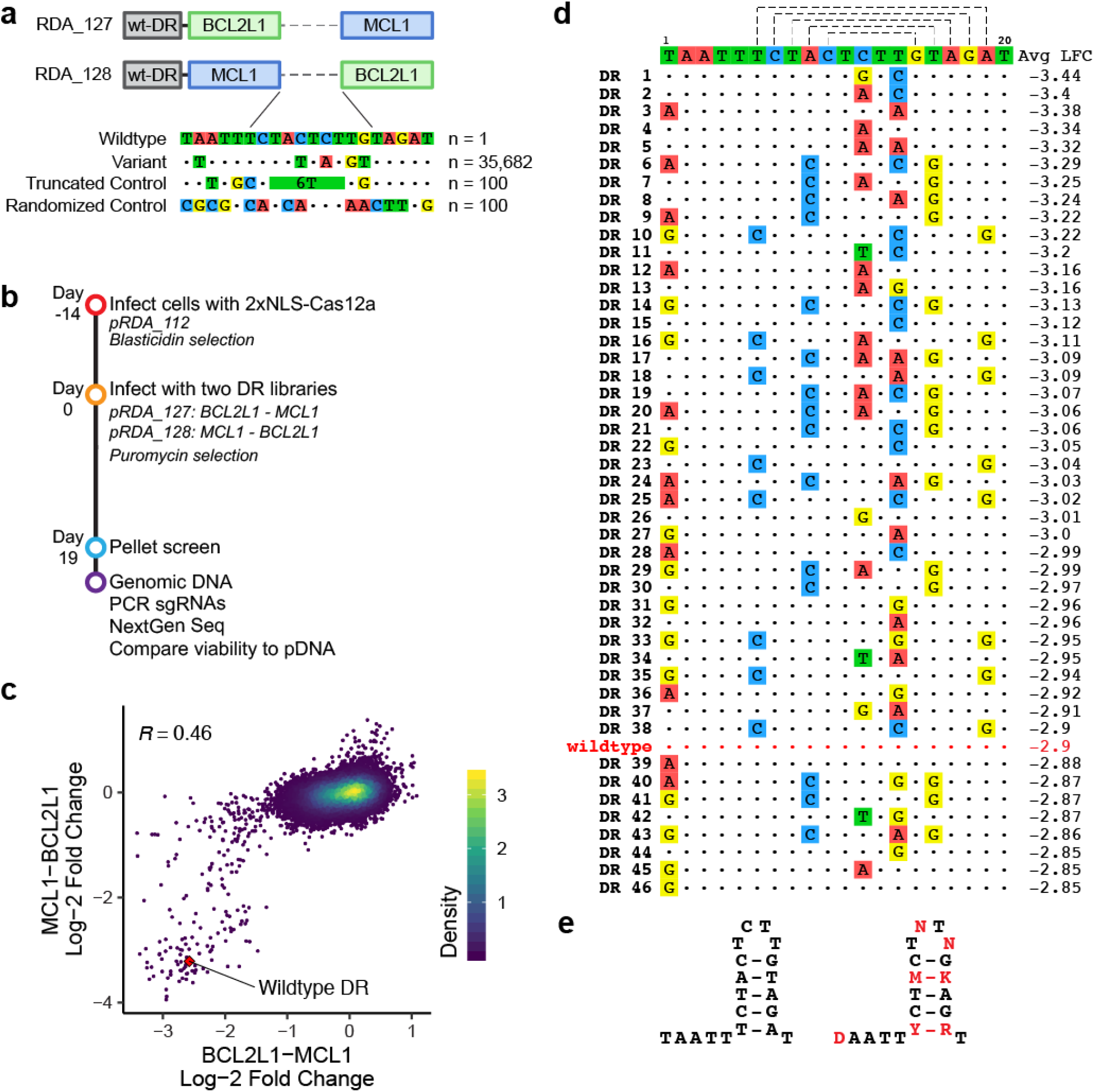
Development of alternate direct repeat sequences for multiplexing with AsCas12a. (**a**) Schematic of experimental design. (**b**) Timeline for executing screens to identify active variant direct repeats. (**c**) Average log2-fold change for direct repeats with both orientations of the BCL2L1 and MCL1 guides. Pearson correlation coefficient is indicated. (**d**) Most active variant direct repeat sequences by average log2-fold change across replicates and orientation. The wildtype sequence is on the top line with the nucleotide substitutions for the top variant sequences below. (**e**) Wildtype direct repeat sequence (left) and a consensus sequence for active variant direct repeats (right). Nucleotides in red denote positions that are flexible for alternate sequences.

Examining these active DR variants, we saw clear sequence preferences (**Fig 4d**). First, several single-stranded positions were intolerant of any changes, whereas others tolerated certain nucleotide substitutions. For example, position 1 tolerated A or G but not C, whereas active constructs were observed with all nucleotides at position 12 and 14 in the loop. Interestingly, this observation in the 12th position of the loop agrees with alignment of direct repeat sequences across Cas12a orthologs^33^. All recovered active sequences maintained basepairing in the stem, but basepairing alone was insufficient for activity, as the nucleotide sequence proved important. At the base of the stem, a T-A basepair could be replaced with a C-G basepair, but no other orientations, indicating a preference for a pyrimidine on one side and a purine on the other. Likewise, in the 4th basepair of the stem, only A-T and C-G basepairs showed activity. Other stem positions, however, did not tolerate any substitutions. Thus, we identified a consensus sequence for active variant DRs (**Fig. 4e**), and numerous examples thereof, which should assist in the creation of multiplexed arrays to minimize repetitive sequences.

### Multiplexing to assay synthetic lethal interactions

To assess the effectiveness of multiplexing across numerous endogenous genes, we designed a library targeting synthetic lethal gene pairs. Here, the compact design of AsCas12a arrays leads to significant advantages when synthesizing and sequencing DNA compared to Cas9-based approaches. For AsCas12a, a second guide requires only 43 additional nucleotides, whereas a second Cas9 guide cassette is 346 nucleotides (**Fig. 5a**).

**Figure 5.**
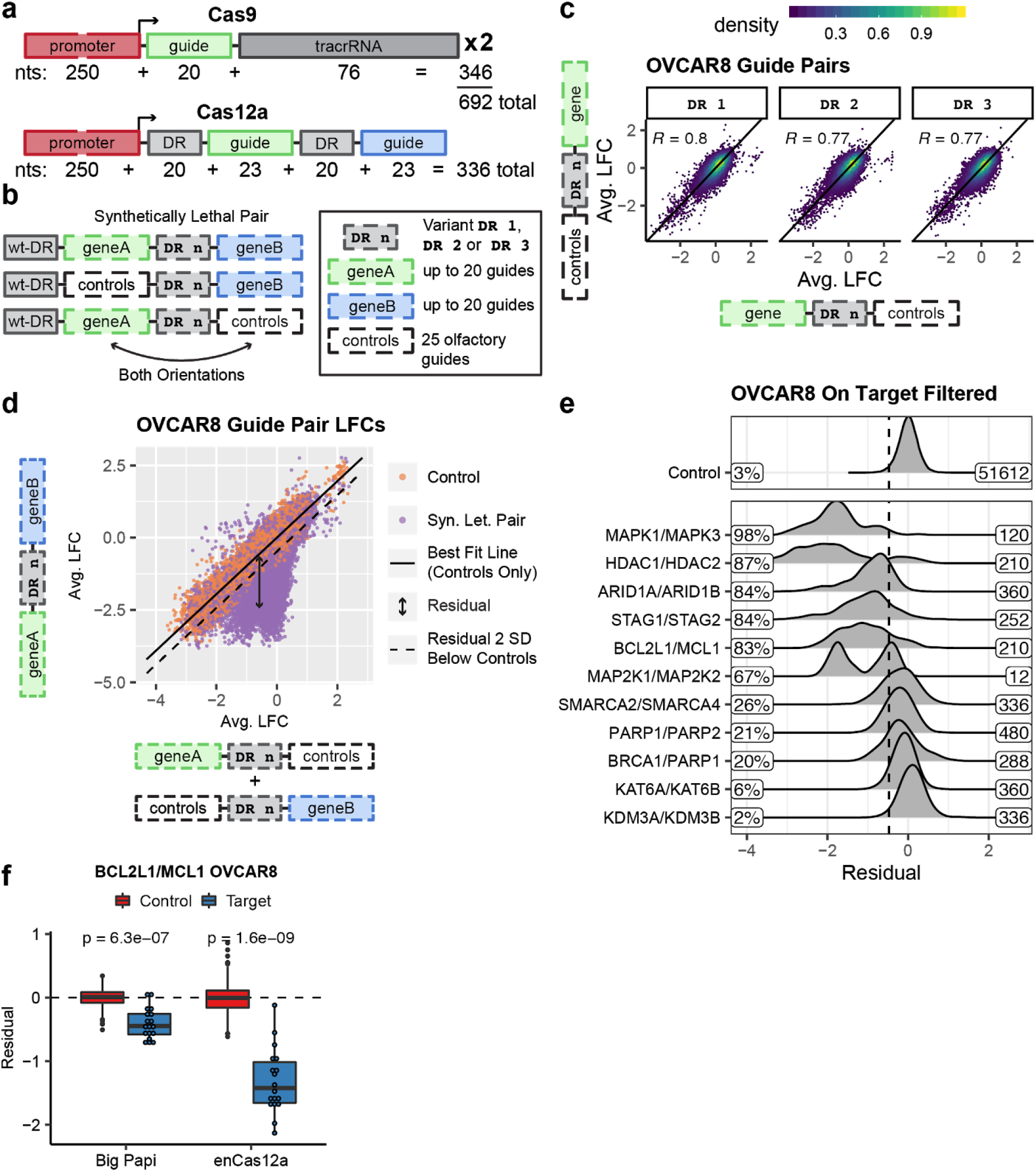
Validation of AsCas12a performance with synthetic lethal gene pairs. (**a**) Comparison of DNA cassettes necessary for dual knockout with Cas9 versus Cas12a. (**b**) Schematic of library design. Numbered direct repeats reference the same sequences as figure 4. (**c**) Correlation between the average log2-fold change (LFC) of target guides in position 1 versus position 2 for all three DR variants screened in OVCAR8. (**d**) Average LFC for guide pairs versus the sum of each guide paired with controls in OVCAR8. Control points represent guide pairs with one control guide and one target guide. Regression line fit with control points only. Dashed line represents a residual two standard deviations below the mean residual for control points. (**e**) Density of residuals for synthetic lethal guide pairs in OVCAR8, filtered for guides with a Seq-DeepCpf1 score greater than 50. Dashed line represents two standard deviations below the mean residual of controls. Percent of pairs with a residual to the left of the dashed line is included. Labeled on the right is the number of unique constructs in the distribution. (**f**) Comparison of residuals for BCL2L1/MCL1 in OVCAR8 across Cas platforms. Libraries were filtered and residuals recalculated to account for differences in library design. Control constructs have one target guide (BCL2L1 or MCL1) and one control guide (n=180), whereas target constructs contain a synthetic lethal guide pair. P-values were calculated using a one-sided t-test with the alternative hypothesis that the mean of the target population was less than the mean of controls. Boxes represent the 25th, 50th and 75th percentiles, whiskers show 10th and 90th percentiles.

We targeted 12 gene pairs we and others have previously identified as synthetic lethal^6,22,34^. For each gene in a given pair, we randomly selected up to 20 guides, 3/4 and 1/4 of which targeted TTTV and TTTT PAMs, respectively (**Fig. 5b**). We synthesized guide pairs in both orientations, and separated each with the top three variant DRs described above; each gene pair was thus assessed with up to 2,400 unique constructs (20 guides × 20 guides × 2 orientations × 3 DRs). To account for single guide effects we also paired each guide with 25 guides targeting olfactory receptors as controls. We screened this library in both A375 and OVCAR8 cells engineered to express enCas12a.

Following PCR and sequencing, we calculated the log2-fold-change (LFC) relative to pDNA and saw good correlation between replicates in both cell lines (**Supplementary Fig. 7a**). For the olfactory controls, we observed that two guides targeting OR5W2 and OR13H1 had a viability effect on their own, so we removed these guides for downstream analyses (**Supplementary Fig. 7b**). To evaluate how position affects guide activity we compared the average LFCs of the ordered target-control constructs with the reverse orientation control-target constructs. We saw that the LFCs of target guides in position 1 were well correlated with the LFCs in position 2 for all three DR variants in both OVCAR8 and A375 (**Fig. 5c**, **Supplementary Fig. 7c**), suggesting that position has minimal effect on guide activity.

To quantitate synergies between guide pairs we first calculated an expected phenotype by determining the average LFC of each targeting guide when it was coupled with controls (**Supplementary Fig. 8a**), and then summing the average LFC for the two guides. We then fit a line between the expected and observed LFCs using constructs with one control and one target guide (**Fig. 5d**). Finally, we calculated the residual from the fit line for all constructs, where a negative residual indicates a synthetic lethal pair. We saw a large fraction of constructs with residuals two standard deviations below the mean of control-target pairs in both OVCAR8 and A375 cells, indicating synthetic lethal interactions. To assess the efficacy of on-target rules in this experimental setting, we filtered the data for guides with a Seq-DeepCpf1 score greater than 50. We saw an increase in the fraction of synthetic lethal constructs for a majority of gene pairs (**Supplementary Fig. 8b, c, d**), confirming the utility of this algorithm.

For some gene pairs we saw a very high fraction of gene pairs score as synthetic lethal. For example, over 80% of guide pairs for STAG1 - STAG2 and HDAC1 - HDAC2 scored in both OVCAR8 (**Fig. 5e**) and A375 cells (**Supplementary Fig. 8b**). ARID1A - ARID1B and MAP2K1 - MAP2K2 also scored with the majority of guides in both cell lines. We also observed cell line specific differences, such as KMD3A - KDM3B, which scored with 34% of guides in A375 cells, but only 2% in OVCAR8 cells (at a threshold at which 3% of control pairs score). That we saw generally good correspondence between the activity of guides targeting essential genes across cell lines (**Supplementary Fig. 1c**) suggests this may reflect a true biological difference.

To compare this AsCas12a system to previous synthetic lethal screens conducted with the “Big Papi” approach using SaCas9 and SpCas9^22^, we first reanalyzed the latter with the same analytical methodology to account for differences in library design strategy (see Methods). For BCL2L1 - MCL1, a gene pair we previously validated with small molecule inhibitors, the magnitude of the residuals is substantially greater with AsCas12a in OVCAR8 cells (**Fig. 5f**) and comparable in A375 cells (**Supplementary Fig. 9a**). For MAPK1 - MAPK3, 98% of guide pairs scored as synthetic lethal in OVCAR8 cells with AsCas12a, with a greater magnitude of residuals than with the Cas9 approach (**Supplementary Fig. 9a**). In contrast, we do not observe synthetic lethality in A375 cells for this gene pair; this difference is likely due to the strong viability effect caused by loss of MAPK1 alone in A375 cells (**Supplementary Fig. 8a**), and thus combined targeting of MAPK1 and MAPK3 shows a buffering relationship. BRCA1 - PARP1 and PARP1 - PARP2 show milder phenotypes with both approaches. The correlation between the residuals from both approaches, albeit from a limited set of gene pairs, suggests the variability in synergies across cell lines largely reflects biological differences (**Supplementary Fig. 9b**). Overall, these results show that AsCas12a is at least comparable, and in some circumstances substantially more potent, than a Cas9-based approach. Given the considerable ease of generating and sequencing AsCas12a combinatorial libraries, we suspect that this will become the technology of choice for such screens.

### Triple knockout with Cas12a

To assess the capacity of AsCas12a to target three genes, we designed guides against the cell surface markers, CD47, CD63, and B2M, which are highly expressed in A375 cells and which do not show a viability effect upon knockout in the DepMap^35^. Using these guides, we generated 6 multiplexed arrays with all possible guide orientations (**Fig. 6a**). We also included 3 single guide constructs, and one empty vector control, and used flow cytometry (**Supplementary Fig. 10**) to quantify the fraction of cells that were edited or not for all three markers. Irrespective of guide position, we observed that approximately 70% of cells showed editing at all three loci at the three timepoints assayed (**Fig. 6b**, **Supplementary Fig. 11**). To assess the potential viability effect of simultaneously targeting three genes at once, we performed a competition assay with no guide, single guide, and triple guide constructs. In this internally-controlled assay, EGFP labels cells without enCas12a (and thus no dsDNA breaks) and we measured the fraction of enCas12a-positive, EGFP-negative cells over time by flow cytometry (**Supplementary Fig. 12a**). We observed that the fraction of EGFP-negative cells decreased by 20 - 40% over time with single guide constructs relative to an empty vector control (**Supplementary Fig. 12b**), evidence of the cutting toxicity inherent to using wildtype Cas enzymes^36,37^. With the triple knockout constructs we saw an additional 10% decrease in viability compared to single knockout, suggesting that the marginal impact of additional dsDNA breaks, at least in this cell type, is minor. Nuclease-deactivated versions of AsCas12a fused to KRAB have been described in mammalian cells^38^, as have transcriptional activators^13,15,38,39^, and such CRISPRa and CRISPRi based approaches will be useful in models where toxicity due to numerous dsDNA breaks is overly-confounding.

**Figure 6.**
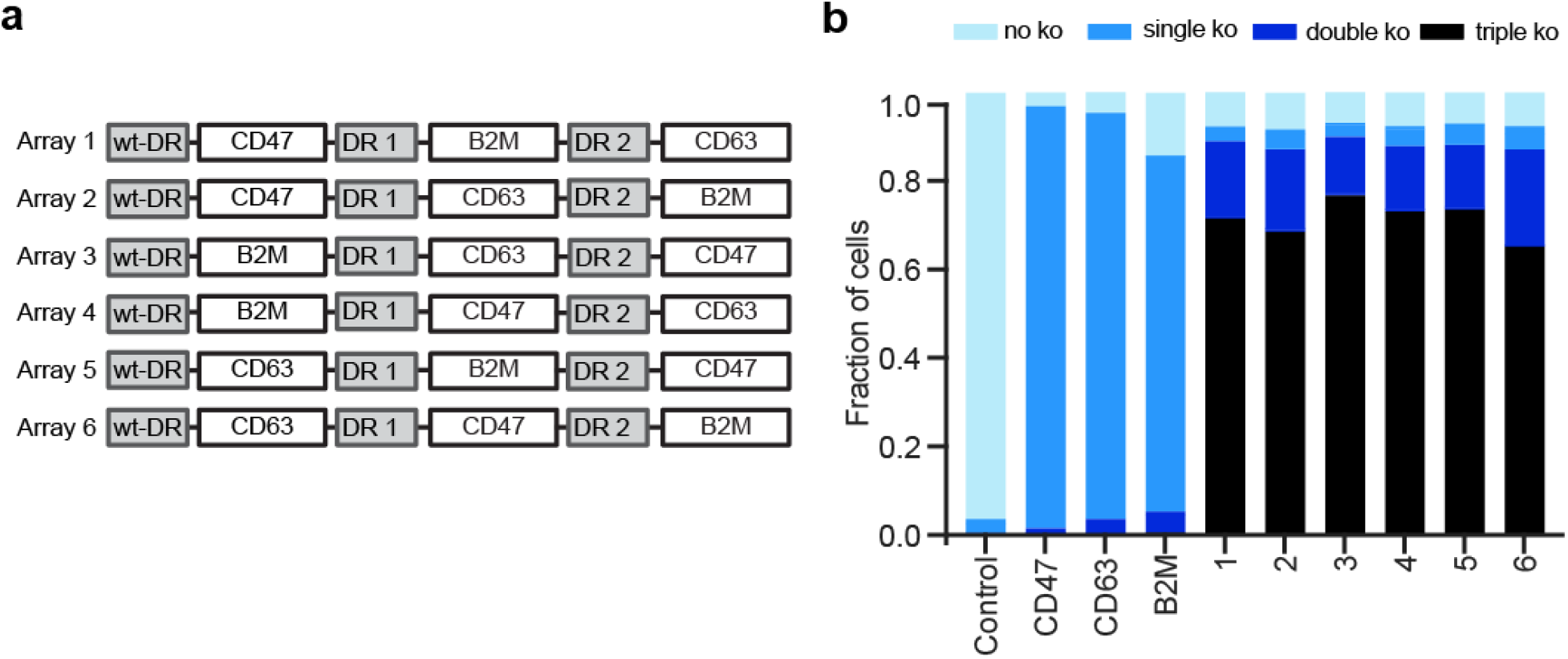
Robust triple knockout with AsCas12a. (**a**) Schematic of 6 multiplexed arrays with guides targeting CD47, B2M and CD63. (**b**) Fraction of cells with no, one, two, or three genes knocked out, assayed by flow cytometry; gates were set such that ~1% of cells score as knockout in the control condition.

## DISCUSSION

Here we present the development of AsCas12a for large scale genetic screens in human cells. We show that highly efficient multiplexing, under conditions of single-copy lentiviral integration, is obtainable with optimized expression constructs, direct repeat sequences, and guide selection rules. We validate this set of optimizations and conduct proof-of-principle combinatorial screens by deeply examining a set of synthetic lethal gene pairs with up to 2,400 unique constructs per pair. We further test this technology by demonstrating triple knockouts in a majority of targeted cells.

The use of enCas12a proved critical to these efforts, both for its improved on-target activity and increased density of available PAM sequences. We validate the Seq-DeepCpf1 on-target scoring algorithm, show that it can be extended to enCas12a, and further show how the input can be modified to score guides with non-canonical PAMs. Additionally, we screened a library of tens of thousands of mismatched guides to develop off-target profiles for AsCas12a constructs, which will enable selection of specific guides. The combination of on- and off-target selection rules for AsCas12a will be available via the GPP sgRNA Designer, broad.io/gpp-sgrna-design.

With an optimized AsCas12a in hand, one clear use-case is combinatorial screens to study genetic interactions. To conduct such studies in high throughput in human cells, researchers have leveraged RNA interference^40^, and more recently CRISPR-Cas9 technologies^6,22,34,41,42^, with the largest GI map to date comprising pairwise interactions among 472 genes^43^, representing 0.05% of all possible interactions of protein-coding genes in the human genome. The massive landscape to be explored, with cell type as an added axis of complexity, recommends the simplicity of synthesizing and sequencing AsCas12a guide constructs compared to Cas9-based approaches, and suggests that the former will become the preferred tool for these studies. Further, the ability to multiplex at higher order will be particularly useful to study gene paralogs, where targeting only one may not reveal a phenotype. Likewise, targeting the same gene with multiple guides in the same construct, which potentially allows the use of compact libraries with only one reagent per gene, has shown some initial promise^44^, and incorporation of the innovations described here makes this an attractive screening approach, especially when cell numbers are limiting. In sum, the results presented here help to establish AsCas12a as a top choice for many applications of genetic screens in human cells.

## ACKNOWLEDGEMENTS

We thank Amy Goodale, Briana Fritchman, and Xiaoping Yang for producing guide libraries and lentivirus; Olivia Bare and Yenarae Lee for logistics support; Caroline Petersen for assistance with screens; Matthew Greene, Adam Brown, Doug Alan, Mark Tomko, and Tom Green for software engineering support; the Broad Institute Genomics Platform Walk-up Sequencing group for Illumina sequencing; the Broad Institute Flow Cytometry Facility for assistance with gating strategy; and the Functional Genomics Consortium for funding support. NIH grant K99-CA218870 to BPK.

## AUTHOR CONTRIBUTIONS

Conceived of the study: XP, AH, JGD

Supervised the project: JGD

Generated 2x-NLS-Cas12a construct: TT, XP

Provided sequence of enAsCas12a in advance of publication: BPK, JKJ

Designed libraries: MH, PCD, REH

Executed genetic screens: KRS, AKS, CS, SMB

Performed analyses: PCD, MH, KRS, JGD

Created visualizations: PCD, AKS, KRS

Curated data: PCD, MH

Wrote the manuscript: KRS, PCD, AKS, REH, JGD

## SUPPLEMENTARY DATA

**Supplementary Data 1**. Reads from tiling libraries for on target performance of 1xNLS-Cas12a, 2xNLS-Cas12a, enCas12a and SpCas9.

**Supplementary Data 2**. Reads from mismatch libraries for off target effects of 2xNLS-Cas12a and enCas12a.

**Supplementary Data 3**. Reads from assays to identify alternate DR sequences.

**Supplementary Data 4**. Reads from the synthetic lethality library screened with enCas12a.

## DATA AVAILABILITY

The read counts for all screening data and subsequent analyses are provided as Supplementary Data and are currently being deposited with the Sequence Read Archive.

## STATISTICAL ANALYSIS

Correlation coefficients were calculated in base R or numpy in Python. Precision recall curves were generated using custom scripts in Python. Wilcoxon rank sum tests for filtering tiling data were performed in R.

## CODE AVAILABILITY

All custom code used for analysis and notebooks are available on GitHub: https://github.com/PeterDeWeirdt

## METHODS

### *Vectors* (Addgene submission in process for some vectors)

Lenti-AsCpf1-Blast: 1x-NLS-Cas12a lentiviral expression construct; also known as pRDA_113; Addgene 84750

pRDA_112: 2x-NLS-Cas12a lentiviral expression construct; also known as pTG_12; Addgene #####

pRDA_174: enCas12a lentiviral expression construct; modified version of pRDA_112 by introduction of point mutations; Addgene #####

pRDA_052: modified version of pLentiGuide for expression of AsCas12a guides; Addgene #####

pRosetta: lentiviral construct for expression of EGFP, puromycin resistance, and blasticidin resistance; Addgene 59700

pRosetta_v2: modification of pRosetta to include a hygromycin resistance cassette; Addgene ######

### Library production

Oligonucleotide pools were synthesized by CustomArray and Twist. BsmBI recognition sites were appended to each guide RNA sequence (whether single guides or tandem guides) along with the appropriate overhang sequences (bold italic) for cloning into the plasmid pRDA_052, as well as primer sites to allow differential amplification of subsets from the same synthesis pool. The final oligonucleotide sequence was thus: 5’-[Forward Primer]CGTCTCA**AGAT**[guide RNA]TTTTTT***GAAT***CGAGACG[Reverse Primer].

Primers were used to amplify individual subpools using 25 μL 2x NEBnext PCR master mix (New England Biolabs), 2 μL of oligonucleotide pool (~40 ng), 5 μL of primer mix at a final concentration of 0.5 μM, and 18 μL water. PCR cycling conditions: 30 seconds at 98°C, 30 seconds at 53°C, 30 seconds at 72°C, for 24 cycles. In cases where a library was divided into subsets unique primers could be used for amplification:

Primer Set; Forward Primer, 5’ – 3’; Reverse Primer, 5’ – 3’

1. AGGCACTTGCTCGTACGACG; ATGTGGGCCCGGCACCTTAA
2. GTGTAACCCGTAGGGCACCT; GTCGAGAGCAGTCCTTCGAC
3. CAGCGCCAATGGGCTTTCGA; AGCCGCTTAAGAGCCTGTCG
4. CTACAGGTACCGGTCCTGAG; GTACCTAGCGTGACGATCCG
5. CATGTTGCCCTGAGGCACAG; CCGTTAGGTCCCGAAAGGCT
6. GGTCGTCGCATCACAATGCG; TCTCGAGCGCCAATGTGACG

The resulting amplicons were PCR-purified (Qiagen) and cloned into the library vector via Golden Gate cloning with Esp3I (Fisher Scientific) and T7 ligase (Epizyme); the library vector was pre-digested with BsmBI (New England Biolabs). The ligation product was isopropanol precipitated and electroporated into Stbl4 electrocompetent cells (Life Technologies) and grown at 30°C for 16 hours on agar with 100 μg mL^−1^ carbenicillin. Colonies were scraped and plasmid DNA (pDNA) was prepared (HiSpeed Plasmid Maxi, Qiagen). To confirm library representation and distribution, the pDNA was sequenced.

### Lentivirus production

For small-scale virus production, the following procedure was used: 24 h before transfection, HEK293T cells were seeded in 6-well dishes at a density of 1.5 × 10^6^ cells per well in 2 mL of DMEM + 10% FBS. Transfection was performed using TransIT-LT1 (Mirus) transfection reagent according to the manufacturer’s protocol. Briefly, one solution of Opti-MEM (Corning, 66.25 μL) and LT1 (8.75 μL) was combined with a DNA mixture of the packaging plasmid pCMV_VSVG (Addgene 8454, 250 ng), psPAX2 (Addgene 12260, 1,250 ng), and the transfer vector (e.g., pLentiGuide, 1,250 ng). The solutions were incubated at room temperature for 20–30 min, during which time media was changed on the HEK293T cells. After this incubation, the transfection mixture was added dropwise to the surface of the HEK293T cells, and the plates were centrifuged at 1,000g for 30 min at room temperature. Following centrifugation, plates were transferred to a 37 °C incubator for 6–8 h, after which the media was removed and replaced with DMEM + 10% FBS media supplemented with 1% BSA and 1% penicillin/streptomycin. Virus was harvested 36 h after this media change.

A larger-scale procedure was used for pooled library production. 20-24 h before transfection, 1.8 × 10^7^ HEK293T cells were seeded in a 175 cm^2^ tissue culture flask. The transfection was performed similarly to the small-scale production; 6 mL of Opti-MEM, 305 μL of LT1, and a DNA mixture of pCMV_VSVG (5 μg), psPAX2 (50 μg), and 40 μg of the transfer vector were used per reaction. Flasks were transferred to a 37 °C incubator for 6–8 h; after this, the media was aspirated and replaced with BSA-supplemented media. Virus was harvested 36 h after this media change.

### Cell culture

A375, OVCAR8, MelJuSo, 786O, and A549 cells were obtained from the Cancer Cell Line Encyclopedia. HEK293Ts were obtained from ATCC (CRL-3216).

All cell lines were routinely tested for mycoplasma contamination and were maintained without antibiotics except during screens, when the media was supplemented with 1% penicillin/streptomycin. Cell lines were kept in a 37 °C humidity-controlled incubator with 5.0% CO2 and were maintained in exponential phase growth by passaging every 2-4 days. The following media conditions and doses of polybrene, puromycin, and blasticidin, respectively, were used:

A375: RPMI + 10% FBS; 1 μg mL^−1^; 1 μg mL^−1^; 5 μg mL^−1^
HEK293T: DMEM + 10% FBS; N/A; N/A; N/A
Meljuso: RPMI + 10% FBS; 4 μg mL^−1^; 1 μg mL^−1^; 4 μg mL^−1^
OVCAR8: RPMI + 10% FBS; 4 μg mL^−1^; 1 μg mL^−1^; 8 μg mL^−1^
A549: DMEM + 10% FBS; 1 μg mL^−1^; 1.5 μg mL^−1^; 5 μg mL^−1^
786O: RPMI + 10% FBS; 4 μg mL^−1^; 1 μg mL^−1^; 8 μg mL^−1^

Olaparib was obtained from Cayman Chemical Co (10621) and screened at a dose of 500 nM. Talazoparib was obtained from Selleckchem (BMN 673) and screened at a dose of 7.81 nM. S63845 was a gift from Guo Wei and was screened at 250 nM. A-1331852 was obtained from Active Biochem (A-6048) and was screened at a dose of 250 nM. Vemurafenib (S1267) was obtained from Selleckchem and screened at a dose of 2 μM. 6-thioguanine was obtained from Sigma-Aldrich and screened at a dose of 2 μg/mL.

### Determination of antibiotic dose

In order to determine an appropriate antibiotic dose for each cell line, cells were transduced with the pRosetta or pRosetta_v2 lentivirus such that approximately 30% of cells were infected and therefore EGFP+. At least 1 day post-transduction, cells were seeded into 6-well dishes at a range of antibiotic doses (e.g. from 0 μg/mL to 8 μg/mL of puromycin). The rate of antibiotic selection at each dose was then monitored by performing flow cytometry for EGFP+ cells. For each cell line, the antibiotic dose was chosen to be the lowest dose that led to at least 95% EGFP+ cells after antibiotic treatment for 7 days (for puromycin) or 14 days (for blasticidin and hygromycin).

### Determination of lentiviral titer

To determine lentiviral titer for transductions, cell lines were transduced in 12-well plates with a range of virus volumes (e.g. 0, 150, 300, 500, and 800 μL virus) with 3.0 × 10^6^ cells per well in the presence of polybrene. The plates were centrifuged at 640 x g for 2 h and were then transferred to a 37 °C incubator for 4–6 h. Each well was then trypsinized, and an equal number of cells seeded into each of two wells of a 6-well dish. Two days post-transduction, puromycin was added to one well out of the pair. After 5 days, both wells were counted for viability. A viral dose resulting in 30–50% transduction efficiency, corresponding to an MOI of ~0.35–0.70, was used for subsequent library screening.

### Pooled screens

Cell lines stably expressing Cas12a were transduced with guides cloned into the pRDA_052 vector in two biological replicates at a low MOI (~0.5). Transductions were performed with enough cells to achieve a representation of at least 500 cells per guide per replicate, taking into account a 30–50% transduction efficiency. Throughout the screen, cells were split at a density to maintain a representation of at least 500 cells per guide, and cell counts were taken at each passage to monitor growth. Puromycin selection was added 2 days post-transduction and was maintained for 5–7 days. After puromycin selection was complete, each replicate was divided into untreated (i.e. no drug / dropout arms) and small molecule treatment arms, each at a representation of at least 500 cells per guide. 14 days after the initiation of small molecule treatment, cells were pelleted by centrifugation, resuspended in PBS, and frozen promptly for genomic DNA isolation.

### Genomic DNA isolation and sequencing

Genomic DNA (gDNA) was isolated using the Machery Nagel NucleoSpin Blood Maxi (2e7–1e8 cells), Midi (5e6–2e7 cells), or Mini (<5e6 cells) kits as per the manufacturer’s instructions. The gDNA concentrations were quantitated by Qubit. For PCR amplification, gDNA was divided into 100 μL reactions such that each well had at most 10 μg of gDNA. Per 96 well plate, a master mix consisted of 150 μL ExTaq DNA Polymerase (Takara), 1 mL of 10x Ex Taq buffer, 800 μL of dNTP provided with the enzyme, 50 μL of P5 stagger primer mix (stock at 100 μM concentration), and 2 mL water. Each well consisted of 50 μL gDNA plus water, 40 μL PCR master mix, and 10 μL of a uniquely barcoded P7 primer (stock at 5 μM concentration). For future experiments, we recommend the use of Titanium Taq DNA Polymerase (Takara) and the addition of 5% DMSO per well, as we have found that these changes improve PCR efficiency.

PCR cycling conditions: an initial 1 min at 95 °C; followed by 30 s at 94 °C, 30 s at 52.5 °C, 30 s at 72 °C, for 28 cycles; and a final 10 min extension at 72 °C. PCR primers were synthesized at Integrated DNA Technologies (IDT). PCR products were purified with Agencourt AMPure XP SPRI beads according to manufacturer’s instructions (Beckman Coulter, A63880). Samples were sequenced on a HiSeq2500 HighOutput (Illumina) with a custom primer of sequence: 5’-CTTGTGGAAAGGACGAAACACCGGTAATTTCTACTCTTGTAGAT. The first nucleotide sequenced with the primer is the first nucleotide of the guide RNA, which will contain a mix of all four nucleotides, and thus staggered primers are not required to maintain diversity when using this approach. Reads were counted by alignment to a reference file of all possible guide RNAs present in the library. The read was then assigned to a condition (e.g. a well on the PCR plate) on the basis of the 8 nt index included in the P7 primer.

### Screen analysis

Following deconvolution, the resulting matrix of read counts was first normalized to reads per million within each condition by the following formula: read per guide RNA / total reads per condition × 10^6^. Reads per million was then log2-transformed by first adding one to all values, which is necessary in order to take the log of guides with zero reads. For each guide, the log2-fold-change from plasmid DNA (pDNA) was then calculated. All reported log2-fold-changes for dropout screens are relative to pDNA; for positive selection screens with small molecules, the log2-fold-change are calculated relative to the dropout arm (i.e. no small molecule treatment arm).

### On-target modeling

To pick guides we filtered for genes that had a significant spearman correlation across replicates and were more active than control genes (two-tailed Mann-Whitney test). We averaged log2-fold changes for all guides targeting the same loci in selected conditions.

Modeling was done as previously with Cas9^22,23^ with a few additional features. We added a tetramer (e.g. TTTT) feature. We also added physiochemical features, which have proven to be useful^45^. Model hyperparameters were chosen using a random grid search with the Python library *scikit-learn*.

### In-silico mutagenesis

To understand the learned nucleotide features of the on-target models, we started with a random seed sequence, and then for each timestep we randomly changed one nucleotide from the previous step. We restricted sequences to have a TTTN PAM and ensured each new sequence was unique. We iterated sequences over 15,001 timesteps. Then we scored each sequence and took the difference between its score and the score of the sequence from the previous step, yielding 15,000 deltas. Then, for each position and nucleotide, we calculated an average difference for the substitution.

### Calculating expected LFCs

For a construct with the ordered elements guide1, DR, guide2, we say 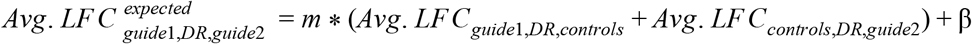, where *m* and β are a fitted slope and intercept. The residual then is the difference between the observed and expected LFCs.

### Filtering SynLet libraries

We used data from the Big Papi SynLet screen in A375 and OVCAR8 cells to compare with the multiplexed enCas12a system. Both libraries targeted the pairs MAPK1/MAPK3, PARP1/PARP2, BCL2L1/MCL1, BCL2L1/BCL2L2, MAP2K1/MAP2K2, and BRCA1/PARP1. We filtered both data sets to account for the heterogeneity of the library designs. After filtering we had 18 observations for each programmed pair (3 guides for each gene in both orientations), and each guide was paired with 15 cutting controls. In the enCas12a library we were left with fewer than 3 guides for MAP2K1, MAP2K2, and BCL2L2 so we removed these genes from the comparison.

To mitigate off-target effects in the enCas12a library we removed guides that were predicted to cut in alternative protein-coding regions between 20% and 100% of the time (Tier I, Bins I and II in the GPP sgRNA design tool). We then tried to maximize on-target efficacy of enCas12a SynLet pairs by removing guides that cut in the first 5% or last 20% of the coding sequence of a gene. Then to match the number of guides in the Big Papi library we used Seq-DeepCpf1 to pick the 3 or 15 best remaining guides for SynLet or control genes respectively.

We filtered the Big Papi library for the same target pairs (3 guides per gene) as well as 15 control guides targeting the cell surface marker CD81 (10 guides) and intronic regions of HPRT1 (5 guides). Note that the original library design for Big Papi already included on and off-target filters.

### Triple knockout with Cas12a

A375 cells stably expressing enCas12a were transduced with 6 triple knockout arrays, 3 single knockout constructs, and one empty control vector. Two days after transduction, cells were selected with puromycin (1μg/mL), and passaged on puromycin for 7 days. Cells were visualized by flow cytometry on the BDAccuri C6 Sampler system 9, 18, and 25 days post infection. To prepare samples for visualization, cells were stained with FITC anti-human CD47 (Biolegend # 323106), PE anti-human CD63 (Biolegend #353004) and APC anti-human β2-microglobulin (Biolegend #316312) antibodies, diluted 1:100 in flow buffer (PBS, 2% FBS, 5μM EDTA), incubated for 30 min on ice, washed with flow buffer twice to remove residual antibody, and resuspended in flow buffer. Flow cytometry data were analyzed using FlowJo (v10). Compensation was applied using single stained empty vector control and triple stained empty vector control cell populations. Zero, single, double, and triple knockout populations were quantified using boolean gating in FlowJo.

