## Supplementary Figures and Tables for "Optimization of AsCas12a for combinatorial genetic screens in human cells"

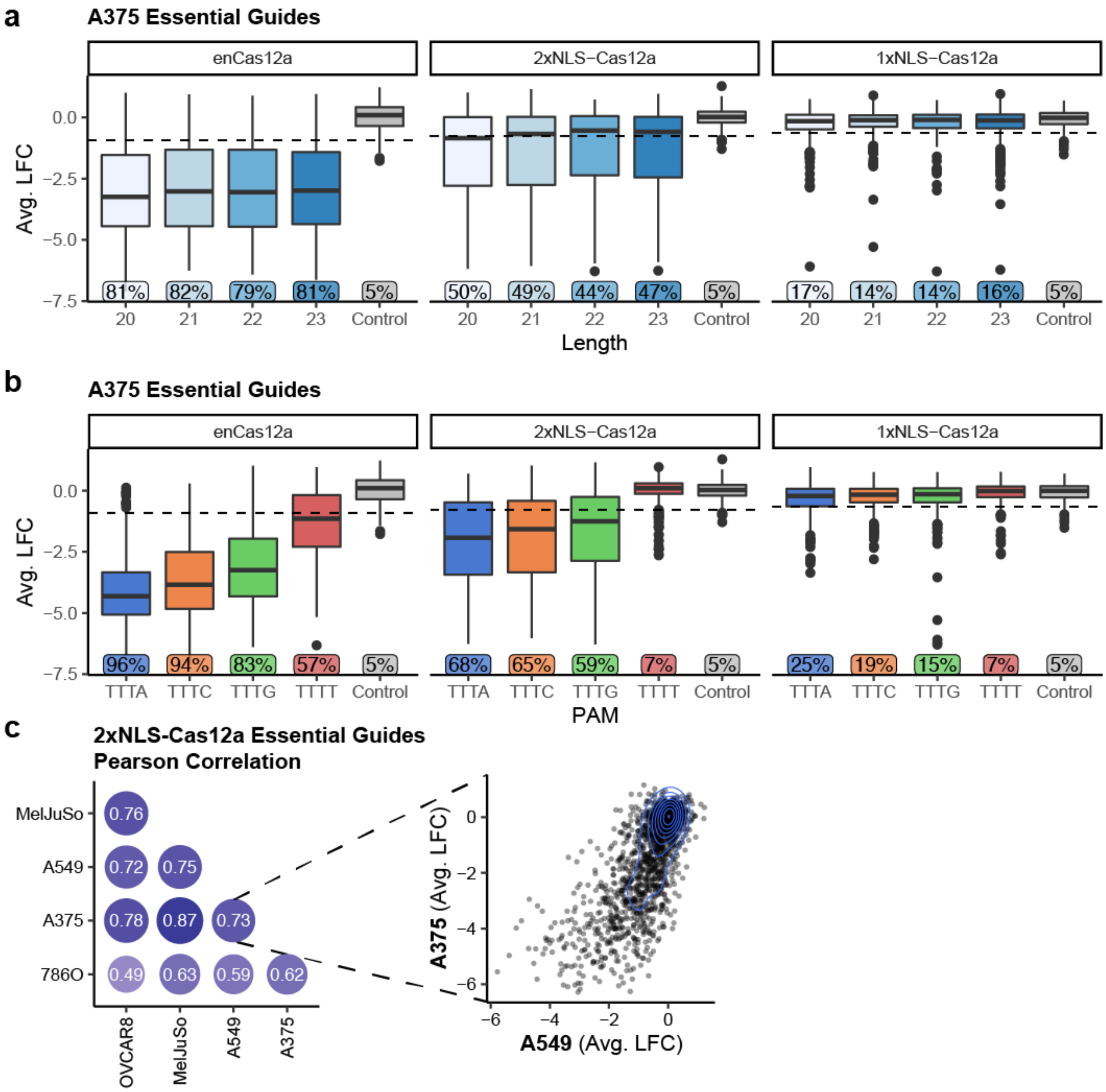


**Supplementary Figure 1** Length and PAM preferences for AsCas12a. (**a**) Activity of guides targeting essential genes (n = 1589) binned by guide length and compared with cell surface control guides (n = 231). Dashed line represents the 5th percentile of control guides. The box represents the 25th, 50th and 75th percentiles, whiskers show 10th and 90th percentiles. (**b**) Same as (a) but binned by PAM. (**c**) Correlation between all essential guides tiled across cell lines with 2xNLS-Cas12a. Size and darkness of each circle corresponds to the indicated pearson correlation coefficient. An example scatter plot comparing guide activity in A375 and A549 cells is shown to the right. Contours represent density of points.

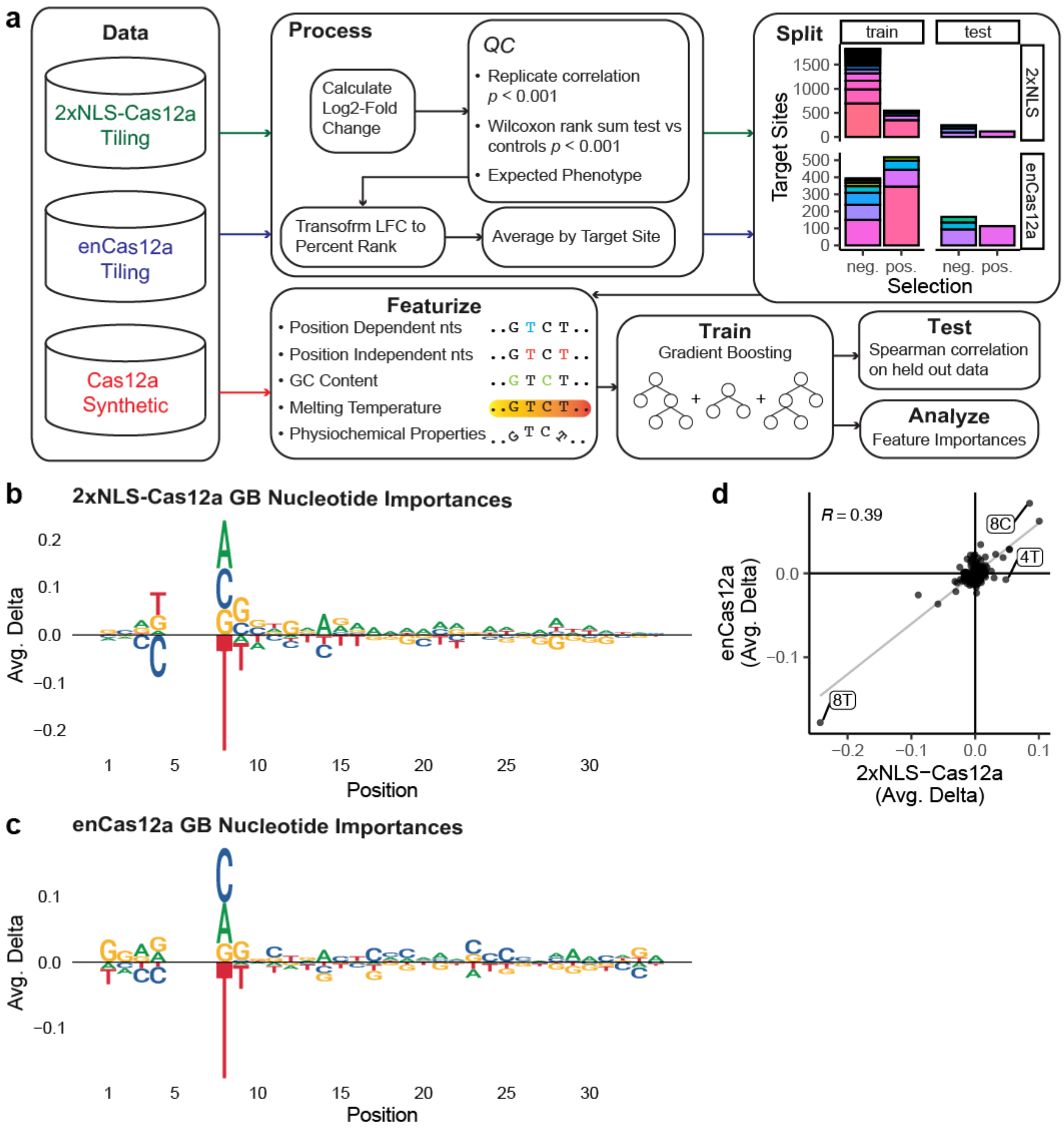


**Supplementary Figure 2** Machine learning to determine on target preferences for AsCas12a. (**a**) Pipeline for gradient boosted models. Colors in the box labelled "split" represent different genes. (**b**) Nucleotide importances for 2xNLS-Cas12a GB from *in-silico* saliency analysis. All tested sequences had a TTTN PAM at positions 5-8. (**c**) Sams as (a) but for enCas12a GB. (**d**) Comparison of nucleotide importances between enCas12a GB and 2xNLS-Cas12a GB. Grey line represents best fit. Labelled points have the largest residual from this line. Spearman correlation is included.

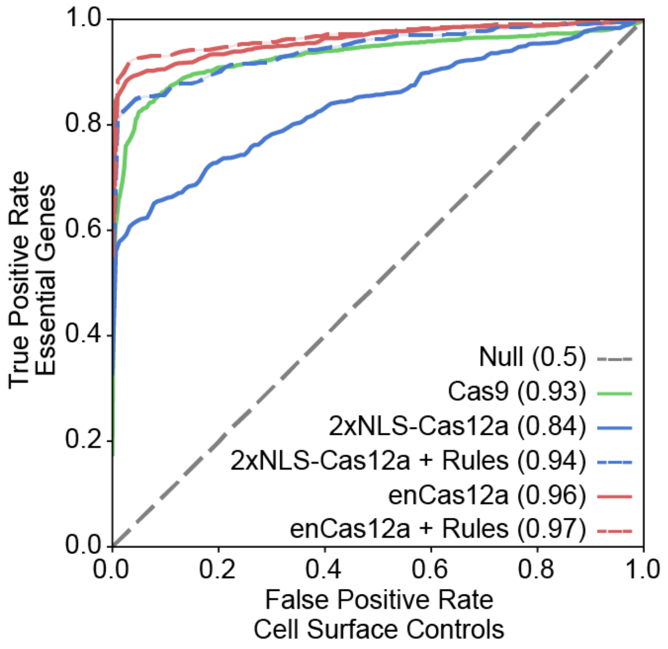


**Supplementary Figure 3** Precision recall curves between essential and cell surface control guides for AsCas12a constructs and SpCas9. Solid lines are the same as in **Fig. 1c**; dashed lines indicate filtering the data for the top half of guides as scored by Seq-DeepCpf1. An area under the curve for each condition is noted in parentheses.

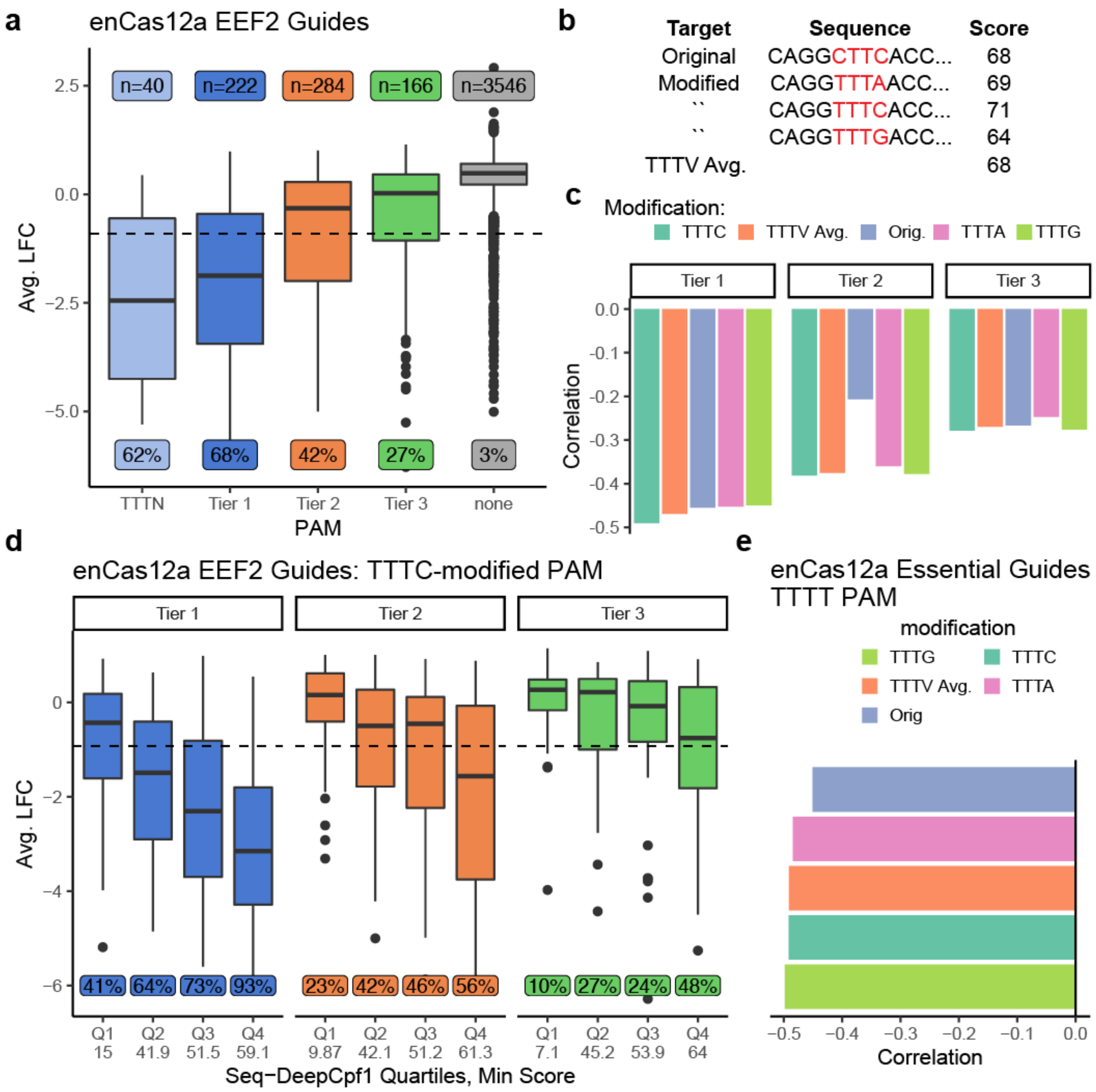


**Supplementary Figure 4** On-target activity scores for non-canonical PAMs with enCas12a. (**a**) Activity of EEF2 guides in A375. Guides are grouped by tier of PAM. Dashed line represents the 5th percentile of flow controls. Fraction of guides beneath this cutoff is included as well as the number of guides in each box. Boxes represent the 25th, 50th and 75th percentiles, whiskers show 10th and 90th percentiles. (**b**) Example of *in-silico* PAM modification. (**c**) Pearson correlation between Seq-DeepCpf1 score and measured log2-fold-change for active PAM tiers in A375 cells. (**d**) Activity of guides in A375 cells binned by TTTC-modified predicted quartile for active PAM tiers. Boxes represent the 25th, 50th and 75th percentiles, whiskers show 10th and 90th percentiles. Minimum score for each quartile is indicated. (**e**) Same as (c), but for TTTT PAM sites.

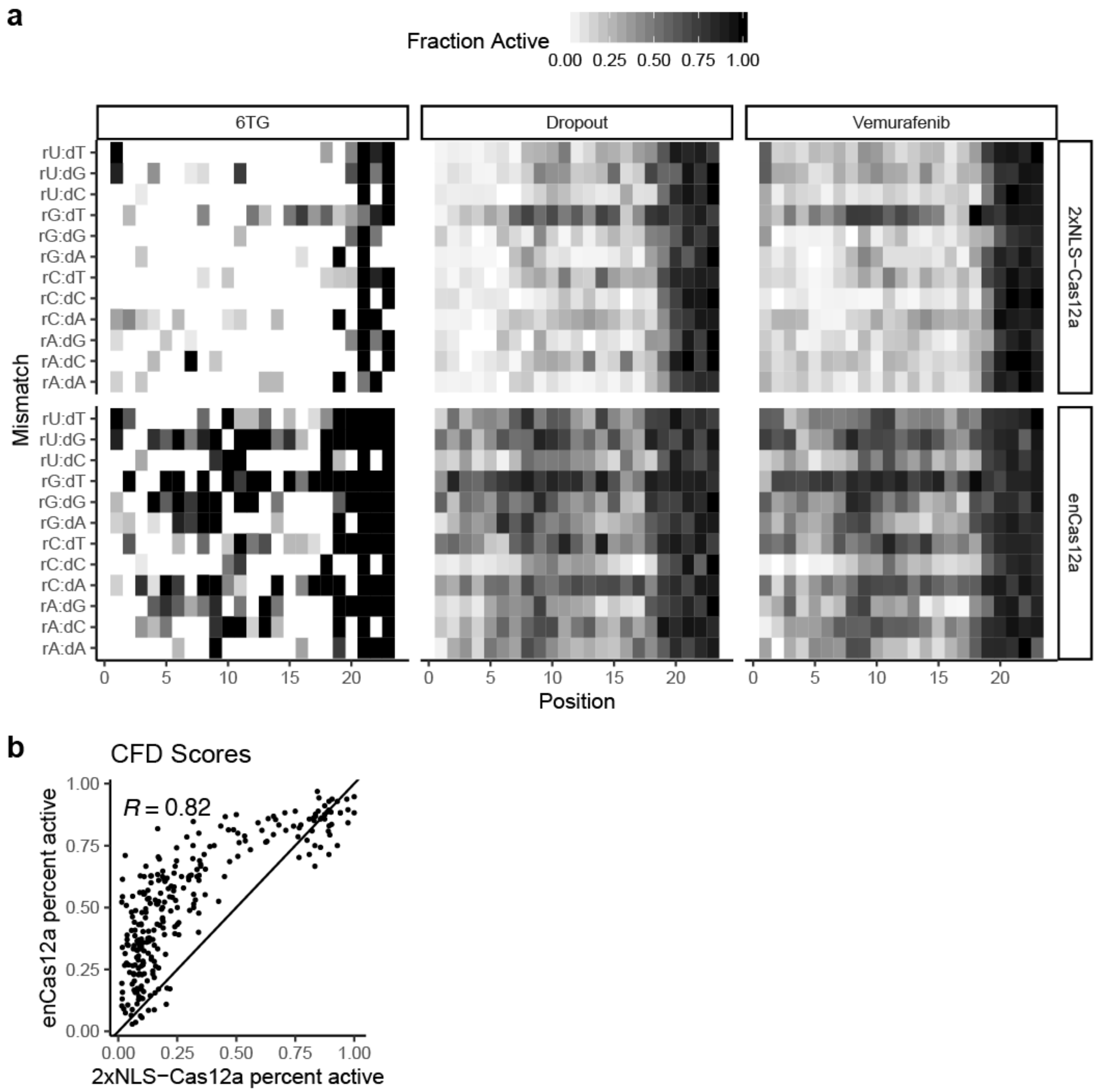


**Supplementary Figure 5** Comparison of off-target profiles. (**a**) Heatmap of the fraction of active guides for each mismatch type and guide position for enCas12a and 2xNLS-Cas12a across three assays. Note that the 6-thioguanine (6TG) conditions has substantially fewer guides than the dropout or vemurafenib screens. Guide position is numbered from PAM proximal to PAM distal. (**b**) Comparison of fraction active for each mismatch/position between enCas12a and 2xNLS-Cas12a. Each point represents one square from the matrices in **Fig. 3c**. Spearman correlation is indicated.

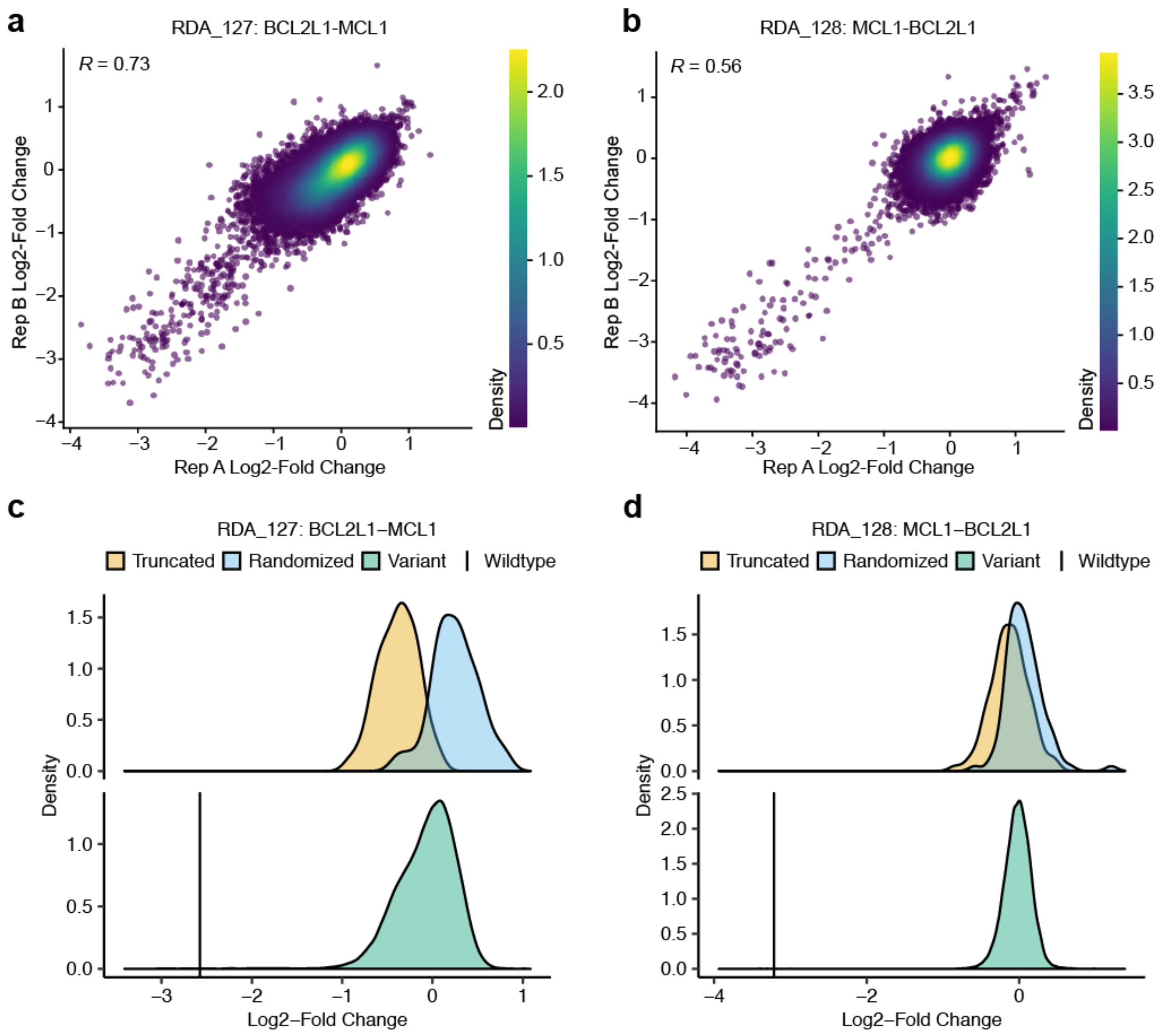


**Supplementary Figure 6** Replicate correlation and control distributions for testing alternate direct repeats. (**a**) Correlation between replicate log2-fold changes for RDA 127, which has the ordered elements: promoter, BCL2L1 guide, direct repeat library, MCL1 guide. Pearson correlation is indicated. (**b**) Same as (a) but for RDA 128 with the ordered elements: promoter, MCL1 guide, direct repeat library, BCL2L1 guide. (**c**) Distribution of log2-fold changes for each type of direct repeat for RDA 127. The two control distributions are on top, whereas variant and wildtype are on bottom. (**d**) Same as (c) but for RDA 128.

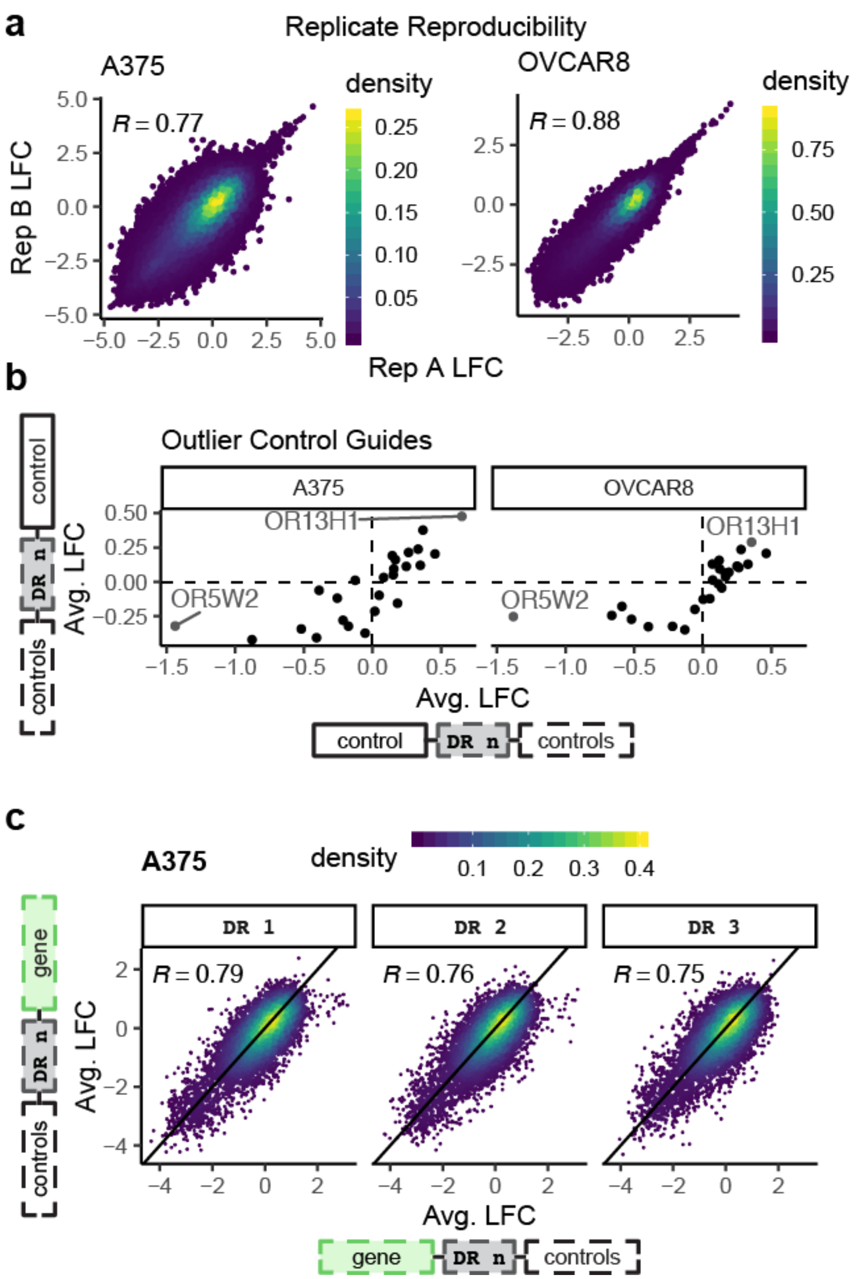


**Supplementary Figure 7** Quality control for synthetic lethality screen. (**a**) Correlation between replicate log2-fold changes in A375 and OVCAR8 cells. Pearson correlation is indicated. (**b**) Average log2-fold changes for control guides. Each point represents a single control paired with all other controls. Axes show each orientation of the guide. Labeled points were removed in downstream analyses. (**c**) Correlation between the average LFC of target guides in position 1 versus position 2 for all three DR variants in A375.

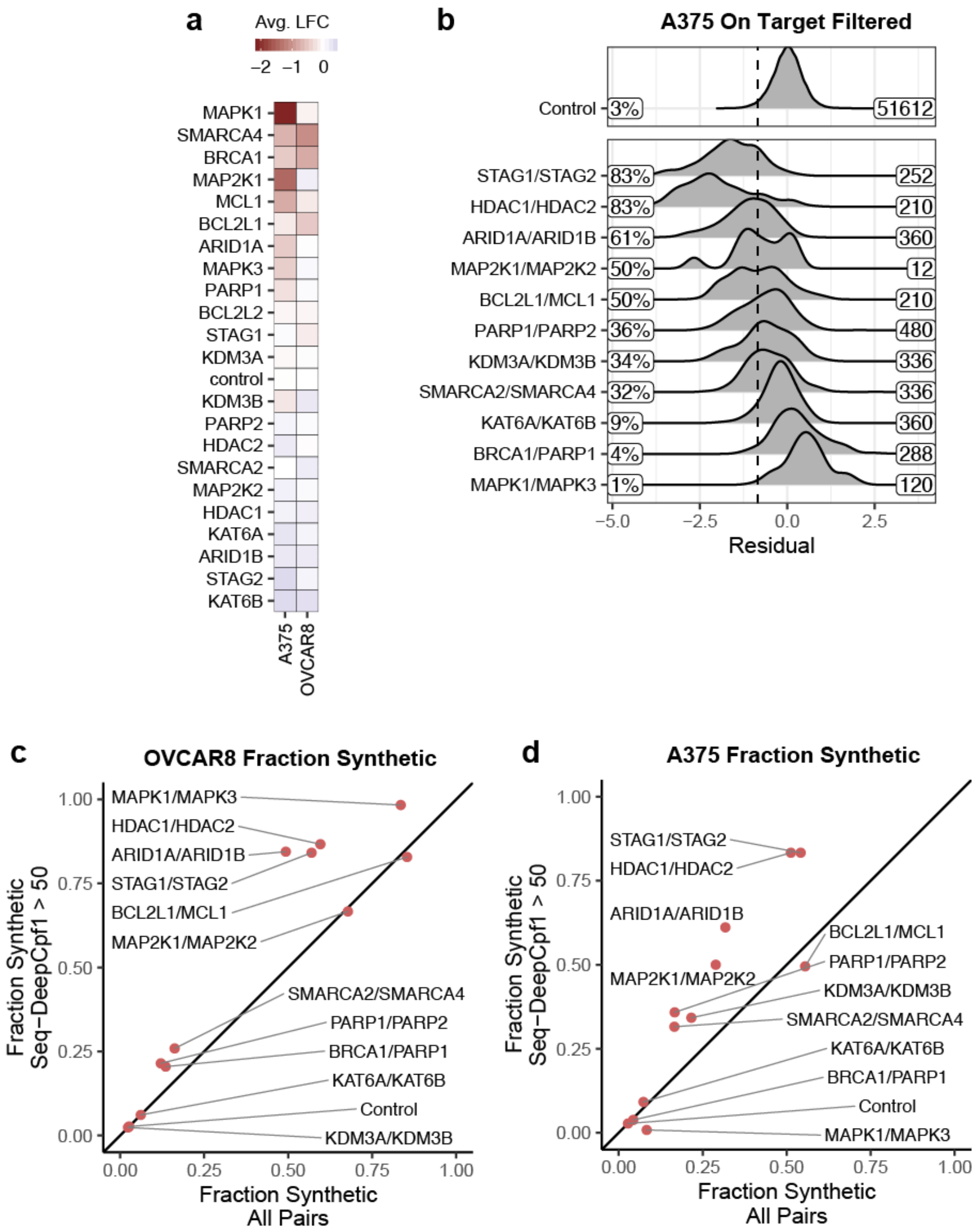


**Supplementary Figure 8** Evaluating synthetic lethal interactions with enCas12a. (**a**) Average log2-fold change for guides paired with controls in A375 and OVCAR8.
(**b**) Density of residuals for synthetic lethal guide pairs in A375, filtered for guides with a Seq-DeepCpf1 score greater than 50. Dashed line represents two standard deviations below the mean residual of controls. Percent of pairs with a residual to the left of the dashed line is included. Labeled on the right is the number of guide pairs in the distribution. (**c**) Comparison of synthetic lethality rates with filtered and unfiltered guides in OVCAR8. (**d**) Same as (b) but in A375.

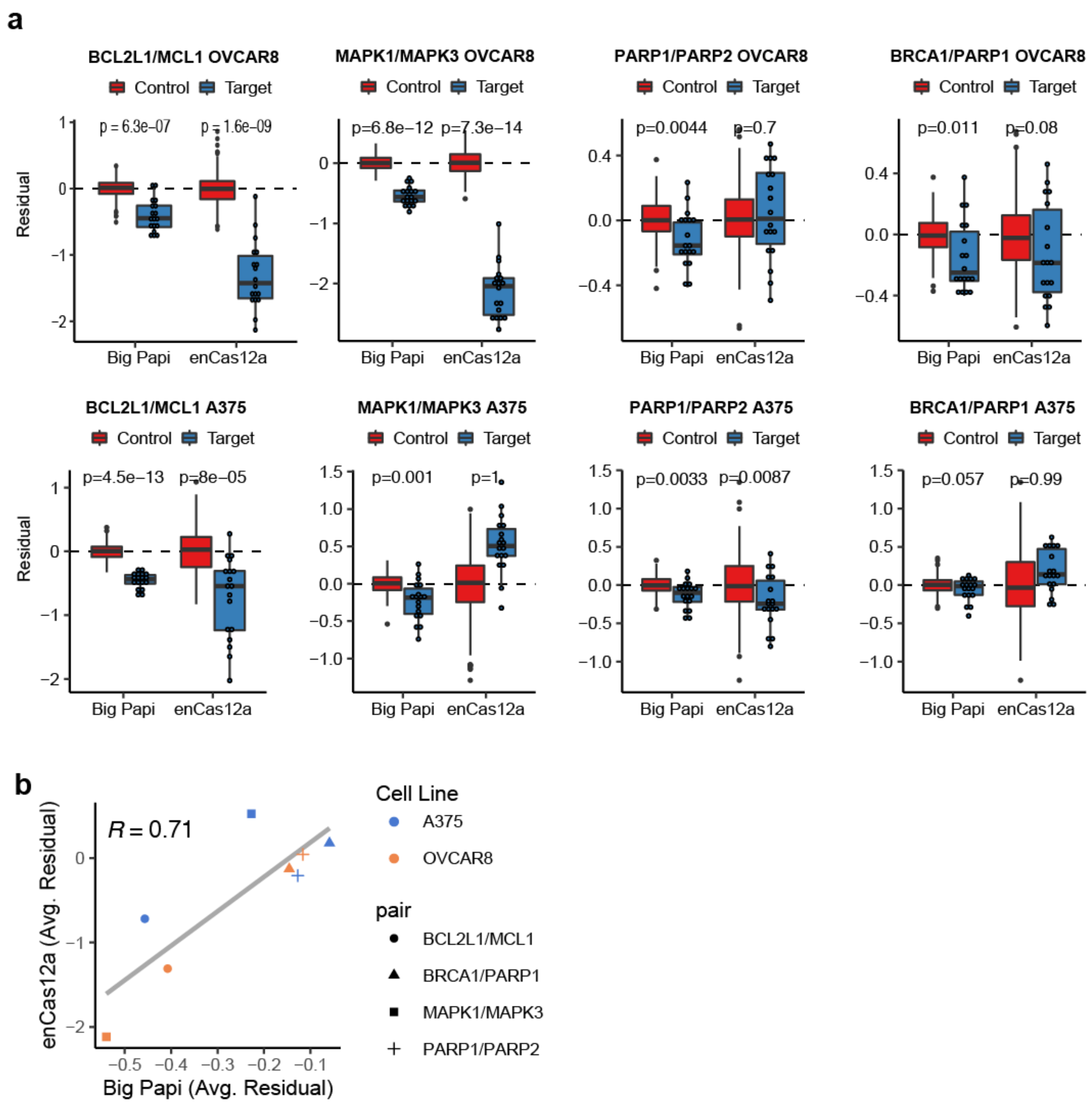


**Supplementary Figure 9** Comparison of synthetic lethality across Cas platforms. (**a**) Residuals for individual synthetic lethal pairs by cell line. Libraries were filtered and residuals recalculated to account for differences in library design strategy. Control constructs have one target guide and one control guide (n=180), whereas target constructs contain a synthetic lethal guide pair. P-value was calculated using a one-sided t-test with the alternative hypothesis that the mean of the target population was less than the mean of controls. Boxes represent the 25th, 50th and 75th percentiles, whiskers show 10th and 90th percentiles. BCL2L1 - MCL1 in OVCAR8 cells repeated from **Fig. 5** for ease of comparison. (**b**) Comparison of the average residual for four synthetic lethal gene pairs using enCas12a or the Big Papi approach. Spearman correlation is included.

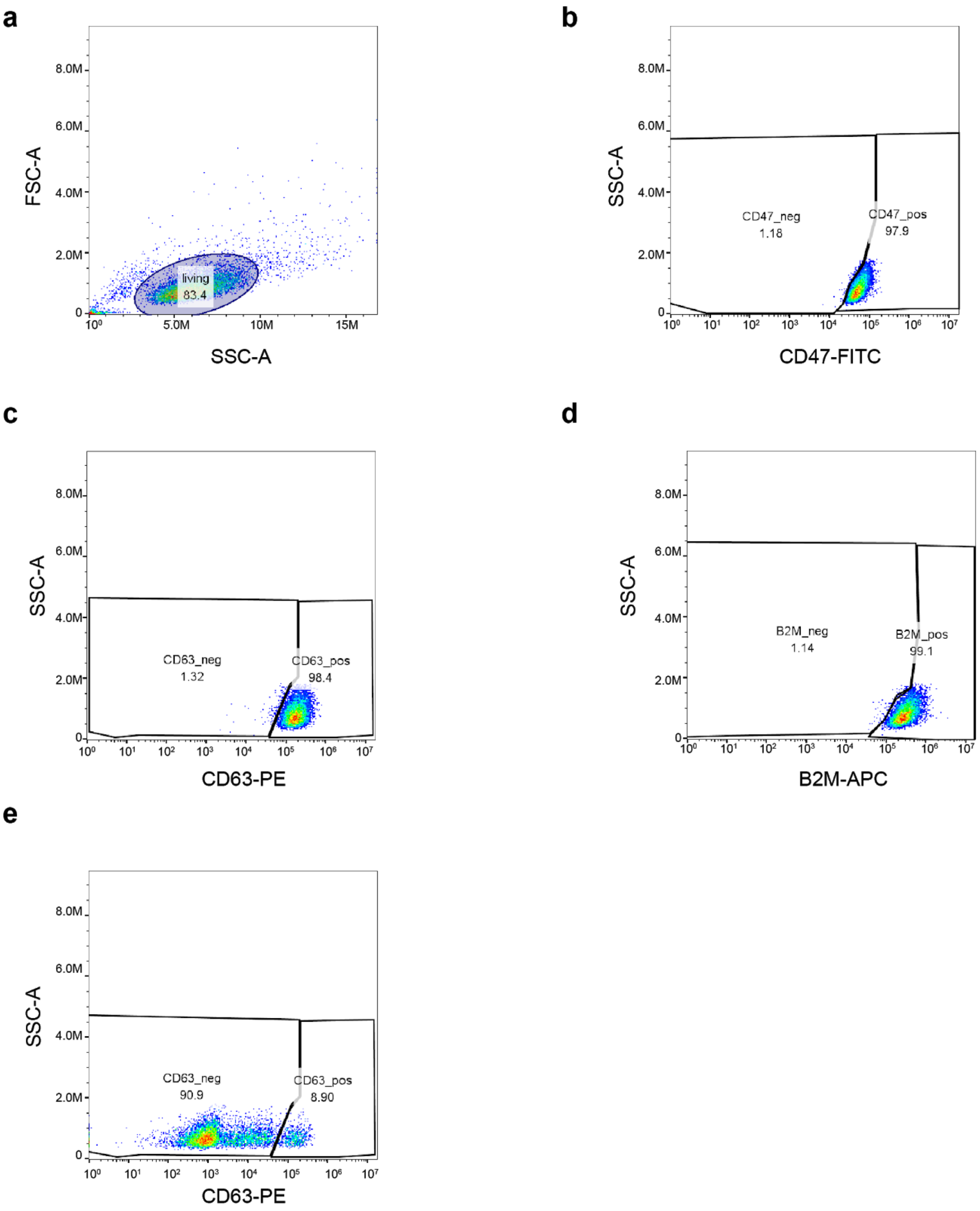


**Supplementary Figure 10** Gating strategy used in Figure 6. (**a**) The live cell population was first gated using forward and side scatter in unstained enCas12a-expressing cells infected with an empty vector control guide. (**b**) CD47 negative gates were set using compensated FL1-A vs side scatter in empty vector control cells stained with all 3 antibodies. (**c**) Same as b, but for CD63 using FL2-A. (**d**) Same as (b) but for B2M using FL4-A. (**e**) CD63 positive and negative populations of representative plot of a representative triple knockout construct (Array 3).

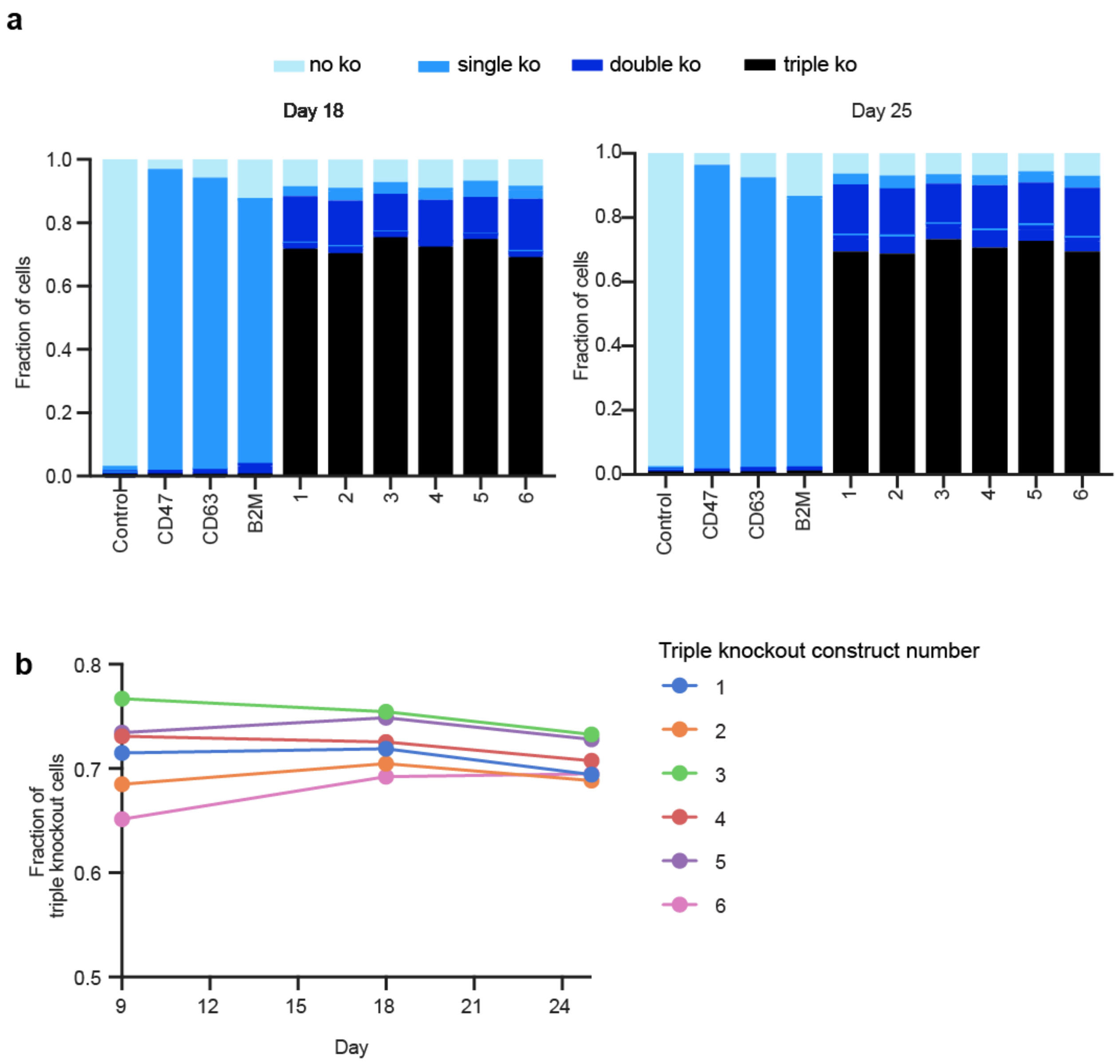


**Supplementary Figure 11** Fraction of edited cells at additional timepoints. (**a**) Fraction of cells with no, one, two, or three genes knocked out, assayed by flow cytometry on day 18 (left) and 25 (right). (**b**) Fraction of cells with triple knockout over time (days 9, 18 and 25.)

**a**

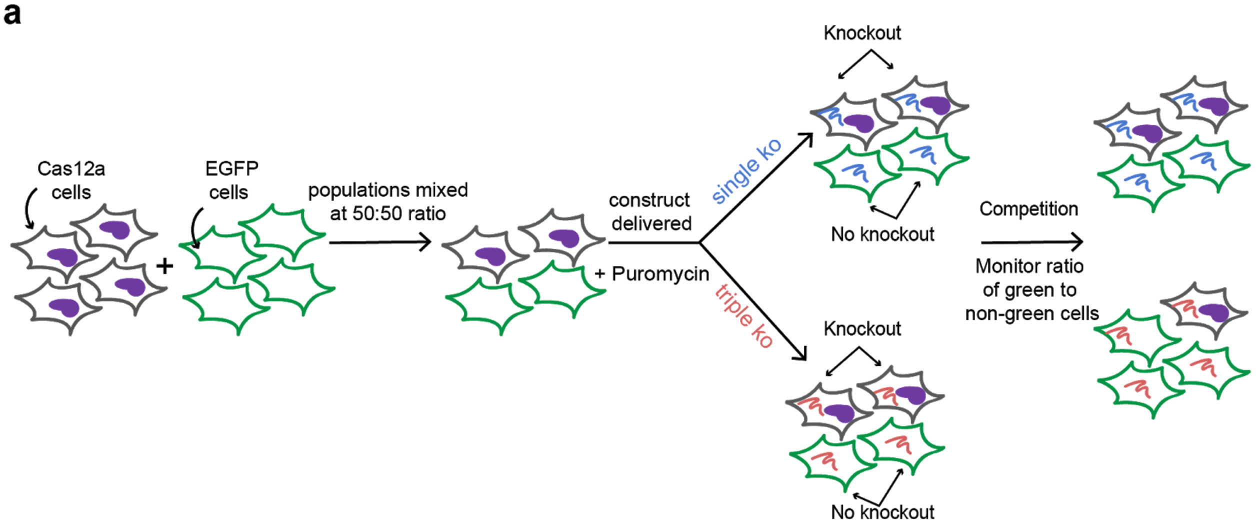


**b**

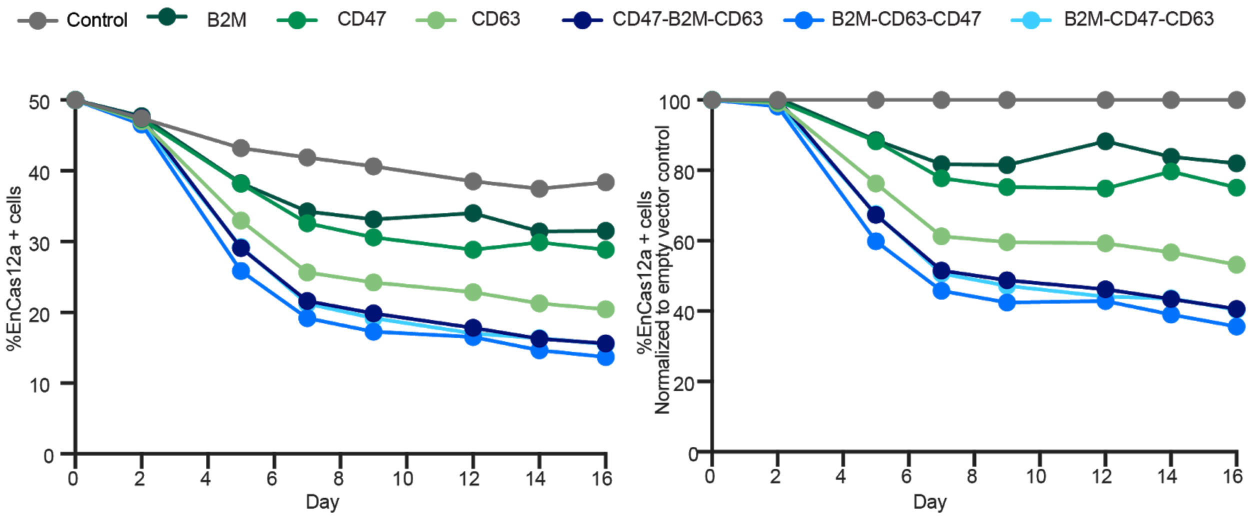


**Supplementary Figure 12** EGFP Competition Assay. (**a**) Schematic of EGFP competition assay to compare viability of no knockout, single knockout and triple knockout of CD47, B2M and CD63. 2 populations of A375 cells, one expressing EnCas12a, the other expressing EGFP, were mixed together at a 50:50 ratio, and were then infected with 3 single knockout constructs, 3 triple knockout constructs, and an empty vector control. Knockout occurs only in EGFP- cells. The fraction of EGFP+ cells to EGFP- cells was assessed over time by flow cytometry (**b**) Fraction of EnCas12a+ cells monitored over time. (**c**) Fraction of EnCas12a+ cells, normalized to the empty vector control.

| Gene ID | # of Guides | Category |
|---|---|---|
| EEF2, HNRNPU, PELP1, TFRC, SF3B1, PSMA6, KPNB1, SNRPD1, RPS20, POLR1C | 1592 | Pan-lethal genes |
| MED12, NF1, NF2, CUL3 | 1640 | Vemurafenib resistance genes |
| HPRT1, NUDT5 | 113 | 6-thioguanine resistance genes |
| BRCA1, BRCA2 | 2628 | Olaparib / Talazoparib sensitive genes |
| BCL2L2, BCL2L1, MCL1, BAX, PMAIP1, BAK1 | 255 | Pro & Anti-apoptotic genes |
| CD81, CD33, FAS, ICAM1 | 231 | Cell surface control genes |
| HRAS, NRAS, PEX6, PEX10, SOX10 | 218 | MelJuSo sensitive genes |
| FBXO42, RNH1, ELOF1, YAP1 | 284 | OVCAR8 sensitive genes |
| TNFSF10, PAX8, STEAP3, SLC25A28, ARNT | 318 | 786O sensitive genes |
| FOXA2, ERBB2, NFE2L2, KRAS, PIK3CA | 941 | A549 sensitive genes |
| EEF2 – Any PAM | 4252 | Guides targeting EEF2 with any PAM |
| **Total Guides:** | 12472 | |

**Supplementary Table 1**. AsCas12a On-target tiling library.

| Condition | Cell Line | | | | |
|---|---|---|---|---|---|
| | A375 | Meljuso | OVCAR8 | 786O | A549 |
| Viability<br>(no small molecule treatment) | Sp: 0.80<br>1x: 0.51<br>2x: 0.86<br>en: 0.80 | 2x: 0.79 | 2x: 0.81 | 2x: 0.42 | 2x: 0.64 |
| S63845 | 1x: 0.32<br>2x: 0.57 | 2x: 0.51 | | | |
| A-1331852 | 1x: 0.51<br>2x: 0.68 | 2x: 0.85 | | | |
| 6-thioguanine | 1x: 0.34<br>2x: 0.88<br>en: 0.90 | | | | |
| Talazoparib | | | 2x:<br>0.17 (250nM)<br>0.15 (7.81nM) | | |
| Olaparib | | | 2x:<br>0.23 (1mM)<br>0.16 (500nM) | | |
| Vemurafenib | 1x: 0.67<br>2x: 0.69<br>en: 0.55 | | | | |

**Supplementary Table 2**. On-target screens with AsCas12a tiling library. 1x: screens with 1xNLS-Cas12a construct; 2x: 2x-NLS-Cas12a construct; en: enCas12a construct (see **Fig. 1**a**).** Sp: SpCas9 screen with an analogous library. Screens were performed in duplicate, and the Pearson correlation of the log2-fold-change is indicated.
